# Structural basis for ligand recognition and G protein-coupling promiscuity of the cholecystokinin A receptor

**DOI:** 10.1101/2021.05.09.443337

**Authors:** Qiufeng Liu, Dehua Yang, Youwen Zhuang, Tristan I. Croll, Xiaoqing Cai, Antao Dai, Xinheng He, Jia Duan, Wanchao Yin, Chenyu Ye, Fulai Zhou, Beili Wu, Qiang Zhao, H. Eric Xu, Ming-Wei Wang, Yi Jiang

**Affiliations:** The CAS Key Laboratory of Receptor Research, Shanghai Institute of Materia Medica, Chinese Academy of Sciences, Shanghai 201203, China; University of Chinese Academy of Sciences, Beijing 100049, China; The National Center for Drug Screening, Shanghai Institute of Materia Medica, Chinese Academy of Sciences, Shanghai 201203, China; Department of Haematology, Cambridge Institute for Medical Research, University of Cambridge, Cambridge CB2 0XY, U.K; School of Pharmacy, Fudan University, Shanghai 201203, China; School of Life Science and Technology, ShanghaiTech University, Shanghai 201210, China; CAS Center for Excellence in Biomacromolecules, Chinese Academy of Sciences, Beijing 100101, China; State Key Laboratory of Drug Research, Shanghai Institute of Materia Medica, Chinese Academy of Sciences, Shanghai 201203, China; School of Basic Medical Sciences, Fudan University, Shanghai 200032, China

## Abstract

Cholecystokinin A receptor (CCK_A_R) belongs to family A G protein-coupled receptors (GPCRs) and regulates nutrient homeostasis upon stimulation by cholecystokinin (CCK). It is an attractive drug target for gastrointestinal and metabolic diseases. One distinguishing feature of CCK_A_R is its ability to interact with sulfated ligand and to couple with divergent G protein subtypes, including G_s_, G_i_, and G_q_. However, the basis for G protein coupling promiscuity and ligand recognition by CCK_A_R remain unknown. Here we present three cryo-electron microscopy (cryo-EM) structures of sulfated CCK-8 activated CCK_A_R in complex with G_s_, G_i_, and G_q_ heterotrimers, respectively. In these three structures, CCK_A_R presents a similar conformation, whereas conformational differences in “wavy hook” of Gα subunits and ICL3 of the receptor serve as determinants in G protein coupling selectivity. These structures together with mutagenesis data provide the framework for understanding the G protein coupling promiscuity by CCK_A_R and uncover the mechanism of receptor recognition by sulfated CCK-8.

---

Cholecystokinin (CCK) is one of the earliest discovered gastrointestinal hormones, participating in gallbladder contraction and pancreatic enzyme secretion. It also acts as a neurotransmitter and is extensively distributed throughout the nervous system ^1^. Selective cleavage of CCK precursor produces a series of bioactive isoforms in different lengths, with CCK-58, −33, −22, and −8 comprising the major peptide fragments in humans. However, the carboxy-terminal octapeptide CCK-8 (DYMGWMDF) is well conserved across species and is the smallest form that retains the full range of biological actions ^2^, mediated by two CCK receptor subtypes (CCK_A_R and CCK_B_R), which are present throughout the CNS and the gut. CCK_A_R is primarily expressed in the alimentary tract, while CCK_B_R is mainly found in the brain and the stomach ^3^. CCK_A_R has a ~500-fold higher affinity to CCK that has a sulfated tyrosine, whereas CCK_B_R discriminates poorly between sulfated and non-sulfated CCK ^4^.

CCK regulates appetite and food intake primarily through CCK_A_R on the vagal afferent neurons ^5–8^, making CCK_A_R an attractive therapeutic target for obesity. However, drug development against CCK_A_R is challenging, partly due to limited efficacy and safety concerns. Although several drug candidates are undergoing clinical trials, none has been approved to date ^9,10^. Extensive efforts were made to elucidate the mechanism of agonism at CCK_A_R through mutagenesis studies based on modeled receptor structures ^11–15^. Nonetheless, the lack of precise structural information largely impedes our understanding of the molecular details regarding ligand recognition and receptor activation, thus the drug discovery targeting CCK_A_R.

Most G protein-coupled receptors (GPCRs) are known to couple with a specific subtype of G proteins to elicit intracellular signal transduction ^16–23^. There are four G protein subtypes, *i.e*., stimulatory G protein (G_s_), inhibitory G proteins (G_i_), G_q_, and G_12/13_, participating in signaling pathways involving cAMP (G_s_ and G_i_), calcium (G_q_), and small G protein (G_12/13_). A number of GPCR-G protein complex structures reported recently reveal that the primary determinants of G protein coupling selectivity reside in the C-terminal α5 helix of Gα subunit and relative outward movement of TM6 ^24,25^. However, CCK_A_R is different from most GPCRs for its ability to couple with several subtypes of G proteins. Activation of CCK_A_R elicits a diversified G protein coupling pattern ^26^: predominantly G_q_ ^27^, but G_s_ ^28^, G_i_ ^27,29^, and G_13_ ^30,31^ all play their roles in CCK_A_R signaling. This unique feature makes CCK_A_R an ideal model to study G protein selectivity and promiscuity (Fig. 1a). Here, we report three cryo-EM structures of sulfated CCK-8 activated CCK_A_R in complex with heterotrimeric G_q_, G_s_, or G_i_ protein, respectively. These structures reveal the unique binding mode in ligand recognition and the structural determinants responsible for G protein selectivity and promiscuity of CCK_A_R.

**Fig. 1.**
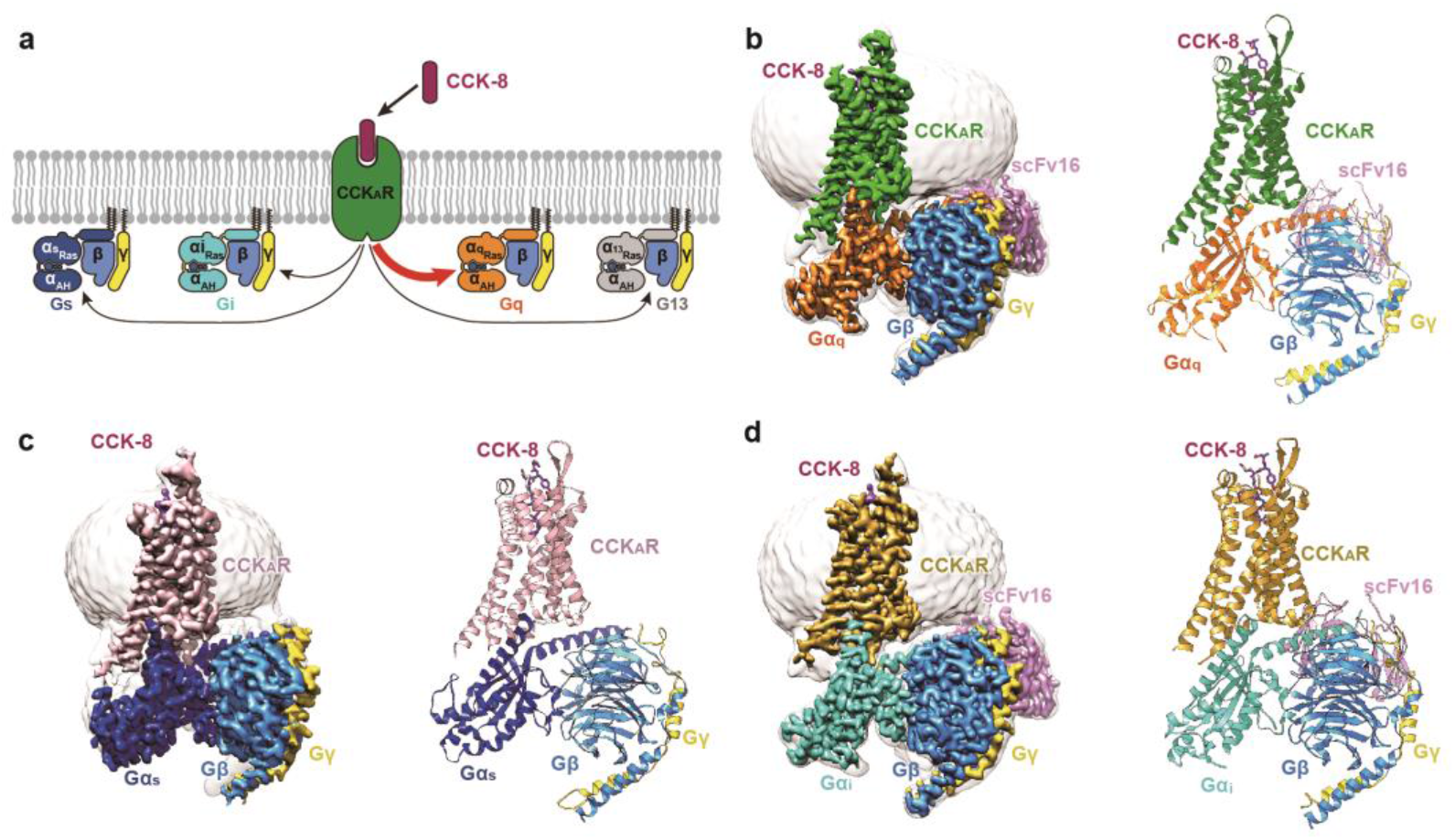
Cryo-EM structures of CCK_A_R–G protein complexes. **a,** Schematic illustration of G protein coupling promiscuity of CCK_A_R. **b-d**, Three-dimensional map (left panel) and the model (right panel) of the CCK-8–CCK_A_R–G_q_–scFv16 (**b**), CCK-8–CCK_A_R–G_s_ (**c**), and CCK-8–CCK_A_R–G_i_–scFv16 (**d**) complexes. CCK-8, magenta; CCK_A_R (**b**), green; CCK_A_R (**c**), pink; CCK_A_R (**d**); dark yellow; Gα_q_, orange; Gα_s_, blue; Gα_i_, cyan; Gβ, light blue; Gγ, yellow; scFv16, light purple.

## Overall structures of CCK_A_R coupled to different G proteins

The structures of sulfated CCK-8 bound CCK_A_R in complex with G_q_, G_s_, or G_i_ heterotrimers were determined by single-particle cryo-EM at a global resolution of 2.9 Å, 3.1 Å, and 3.2 Å, respectively (Fig. 1, Extended Data Fig. 1, Extended Data Table 1). Sulfated CCK-8 (DY^SO3H^MGMWDF-NH_2_), the highest affinity natural ligand of CCK_A_R ^4^, was used to assemble the CCK_A_R–G protein complexes. Three G protein subtypes were engineered to stabilize the CCK_A_R-G protein complexes (Extended Data Fig. 2). Gα_q_ is chimerized by replacing its αN helix with the equivalent region of Gα_i1_ to facilitate scFv16 binding ^32^. Gα_s_ was modified based on mini-Gα_s_ that was used in the crystal structure determination of the G_s_-coupled adenosine A_2A_ receptor (A_2A_R) ^33^. Two dominant-negative (DN) mutations (G203A and A326S ^34^) were introduced to Gα_i1_, and corresponding DN mutations at equivalent sites of Gα_s_ and Gα_q_ were also introduced (Extended Data Fig. 2b). Unless otherwise specified, G_q_, G_s_, and G_i_ refer to respective engineered G proteins, which are used in CCK_A_R structure determination.

The final structures of the CCK-8–CCK_A_R–G protein complexes contain sulfated CCK-8 (residues D^1P^-F^8P^), Gα Ras-like domain, Gβγ subunits, scFv16, and the CCK_A_R residues (E38^N_term^-F385^8.58^, superscripts refer to Ballesteros–Weinstein numbering ^35^). The majority of amino acid side chains, including CCK-8, transmembrane domain (TMD), intracellular loops (ICLs 1-3), and extracellular loops (ECLs 1-3) were well resolved in the final models (Extended Data Fig. 3). Thus, the complex structures provide reliable details to study mechanisms of ligand recognition and G protein coupling.

Globally, CCK_A_R adopts similar overall conformations in all the three structures, with the all-atom root-mean-square deviation (RMSD) at 0.84 for G_q_/G_s_-coupled receptors, and 1.03 for G_q_/G_i_-coupled receptors. The structure of the CCK-8–CCK_A_R–G_q_ complex, which has the highest resolution at 2.9 Å, was used for detailed analysis and mechanistic evaluation of ligand recognition and receptor activation. The inactive and active structures of the closed homolog receptors (inactive: ghrelin receptor, PDB: 6KO5 ^36^; active: neurotensin receptor 1, NTSR1, PDB: 6OS9 ^19^), all belong to the β-branch of the rhodopsin family, are applied for structural comparison. CCK_A_R presents a fully active conformation, resembling the G_i_-coupled NTSR1, displaying a ~9 Å outward movement of TM6 (measured at Cα of residue at position 6.27 in CCK_A_R and ghrelin receptor) and ~4 Å inward shift of TM7 (Cα carbons of Y7.53) compared with the inactive ghrelin receptor (Extended Data Fig. 4, a and b). Similar to the active NTSR1 complex, the conserved residues in “micro-switches” (PIF, ERY, CWxP, and NPxxY) of CCK_A_R display the conserved conformations observed in active GPCRs (Extended Data Fig. 4c).

## Recognition of sulfated cholecystokinin

The sulfated CCK-8 occupies the orthosteric binding pocket comprised of TM3, TM4, TM5-7, and ECL1-3 (Fig. 2, Extended Data Figs. 5, 6), with its C-terminus inserting into the TMD bundle and the N-terminus facing the extracellular vestibule (Fig. 2a). The binding pocket of CCK-8 is largely overlapped with that of other reported endogenous neuropeptides, such as neurotensin (NTS_8-13_, PDB: 6OS9 ^19^), angiotensin II (Ang II, PDB: 6OS0 ^37^), orexin B (OXB, PDB: 7L1U ^38^), and arginine vasopressin (AVP, PDB: 7DW9 ^39^). Noteworthily, the extracellular side of these neuropeptides undergo remarkable conformational shifts, while their intracellular parts converge in an approximately overlapped position at the bottom of the binding pocket (Extended Data Fig. 7).

**Fig. 2.**
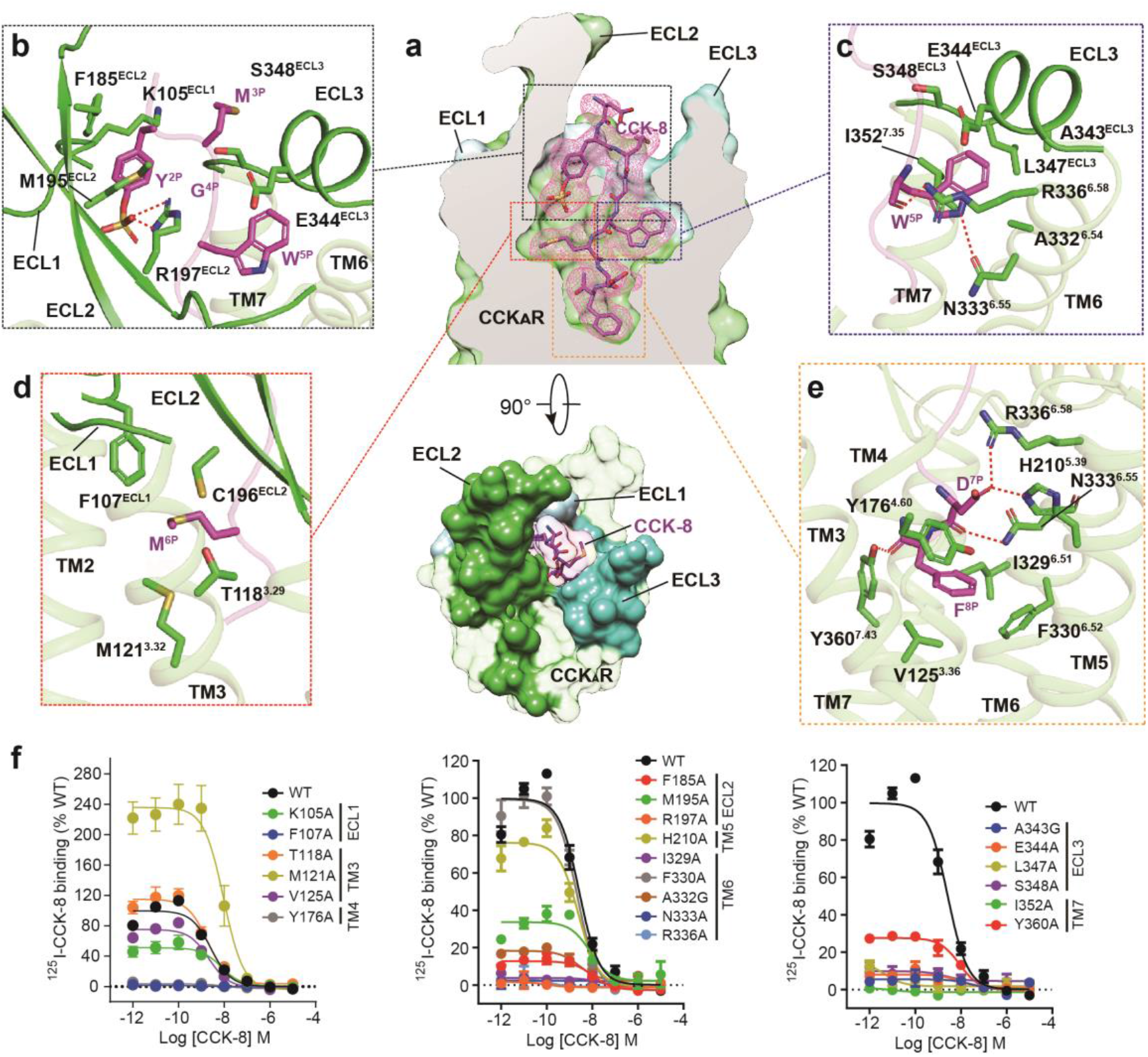
Recognition of sulfated CCK-8 by CCK_A_R. **a**, CCK-8 sits in the orthosteric binding pocket of CCK_A_R shown at side view (upper panel) and extracellular view (bottom panel). The density map of CCK-8 is shown as magenta mesh, and CCK-8 is displayed as magenta sticks. CCK_A_R is shown in green as a cut-away surface (upper panel). ECL1 (light blue), ECL2 (lime green), and ECL3 (turquoise) are highlighted as solid surfaces. **b-e**, Interaction details between sulfated CCK-8 and CCK_A_R. **b**, Recognition of CCK-8 by the three extracellular loops. **c**, Recognition of CCK-8 by the deep hydrophobic cavity beneath ECL3. **d**, Recognition of CCK-8 by the shallow hydrophobic cavity beneath ECL1 and ECL2. **e**, Recognition of CCK-8 by the bottom TMD region. Key interaction residues from CCK_A_R are shown as green sticks, and the receptor is shown in cartoon presentation. Polar interactions are indicated as red dashed lines. **f**, Effects of mutations in the receptor ligand-binding pocket on CCK-8 binding activity assessed by a radiolabeled ligand binding assay (n=3-4). Competition curves of mutants from ECL1, TM3, TM4 (left), ECL2, TM5, TM6 (middle), ECL3 and TM7 (right) compared to wild-type (WT) CCK_A_R are shown.

Of interest is that the octapeptide CCK-8 almost completely occupies the polypeptide-binding pocket, structurally supporting the fact that it is the smallest active form of CCK isoforms. The binding modes of CCK-8 are highly conserved in all three CCK_A_R-G protein complexes (all-atom RMSD 0.71 for CCK-8 in G_q_/G_s_-coupled complexes, and 1.18 for CCK-8 in G_q_/G_i_-coupled complexes), supported by clear EM density maps (Fig. 2a, Extended Data Fig. 3). The ligand recognition region by CCK_A_R can be divided into three major parts: (i) the extracellular loops, (ii) hydrophobic cavities beneath ECLs, and (ii) the bottom of the TMD pocket (Fig. 2a).

At the extracellular side, three ECLs are folded to embrace the N-terminal amino acids of CCK-8 (Fig. 2a). The sulfate group of Y^2P^ ionic interacts with the side chain of R197^ECL2^. This polar interaction prompts the aromatic ring of Y^2P^ to form hydrophobic contacts with F185^ECL2^, M195^ECL2^, and the main chain of K105^ECL1^, thus connecting CCK-8 to ECL1 and ECL2 (Fig. 2b, Extended Data Fig. 6). These structural observations are consistent with the previous finding that the R197^ECL2^M mutation was 1,470-fold less potent than the wild-type (WT) CCK_A_R ^11^. The alanine mutation of R197^ECL2^ completely abolishes the binding of CCK-8, thus strongly supporting the contention that R197^ECL2^ serves as a determinant to discriminate between sulfated and nonsulfated CCK (Fig. 2f, Extended Data Table 2). Likewise, poor ligand selectivity of CCK_B_R may be attributed to a substitution of arginine for valine at the corresponding position (Extended Data Fig. 5). Meanwhile, M^3P^, G^4P^, and W^5P^ clamp the interior surface of ECL3 (Fig. 2b).

Two hydrophobic cavities exist below the ECLs to accommodate M^5P^ and W^6P^ (Fig. 2c, d). The side chain of W^5P^ is sandwiched by the side chains of I352^7.35^ and R336^6.58^ and buries in a deep hydrophobic pocket comprised of TM6, ECL3, and TM7 (Fig. 2c). The backbone CO group of W^5P^ forms an H-bond with R336^6.58^, and its indole nitrogen atom makes another H-bond with N333^6.55^ (Fig. 2c, Extended Data Fig. 6), which is reported to be critical to CCK_A_R activation ^40^. Alanine mutations in residues N333^6.55^, R336^6.58^, A343^ECL3^, E344^ECL3^, L347^ECL3^, and S348^ECL3^ completely abolish the binding of CCK-8, suggesting the key roles of these residues in CCK-8 recognition (Fig. 2f, Extended Data Table 2). In contrast to the W^5P^-occupied hydrophobic pocket, M^6P^ sits in a relatively shallow hydrophobic cavity in the opposite direction, constituted by F107^ECL1^, C196^ECL2^, T118^3.29^, and M121^3.32^ (Fig. 2d, Extended Data Fig. 6). Mutating F107^ECL1^ and residues in ECL2 and ECL3 to alanine eliminated the binding ability of CCK-8 entirely, highlighting an essential function of the three ECLs in peptide recognition (Fig. 2f, Extended Data Table 2).

At the bottom of the binding pocket, D^7P^ and main chain CO group of CCK-8 form a stabilizing polar interaction network with TM5 (H210^5.39^), TM6 (N333 ^6.55^ and R336^6.58^), and TM7 (Y360^7.43^) (Fig. 2e, Extended Data Fig. 6). The phenyl ring of F^8P^ makes polar hydrogen-pi interaction with Y176^4.60^, and inserts into a large hydrophobic crevice comprised of residues from TM3, TM4, TM5, and TM6 (Fig. 2e, Extended Data Fig. 6). Besides N333^6.55^ and R336^6.58^, which also polar interact with W^5P^, I329^6.51^ is closely related to CCK-8 binding (Fig. 2f, Extended Data Table 2).

Elucidation of the recognition mechanism of CCK-8 provides clues for therapeutic development against CCK_A_R. GW-5823, CE-326597, and Glaxo-11p are small molecule agonists for CCK_A_R with moderate activities^10,41,42^. Docking of these agonists to the CCK_A_R shows that they only occupy the bottom half of the TMD binding pocket, thus lacking essential interactions with ECLs1-3 of CCK_A_R (Extended Data Fig. 8). This structural feature may lead to a weaker activity of these small molecule agonists relative to CCK-8. Together, our data provide a framework for understanding the mechanism of small molecule agonist recognition and offer a template for guiding drug design targeting CCK_A_R.

## Overall coupling mode of CCK_A_R-G protein complexes

Although all the four G protein subtypes were reported to interact with CCK_A_R ^26^, only three of the CCK_A_R–G protein samples (CCK_A_R–G_q_, CCK_A_R–G_s_, and CCK_A_R–G_i_ protein complexes) were obtained for high-resolution cryo-EM structure determination (Fig. 1). Structural comparison indicated that TM6 and ICL2 in CCK_A_R adopt nearly identical conformations in G_q_-, G_i_-, and G_s_-coupled structures (Fig. 3a, Extended Data Fig. 7). However, slightly different tilts of the Gα α5 helix were seen among the three heterotrimeric G proteins (4° for Gα_q_/Gα_s_ and 8° for Gα_q_/Gα_i_) (Fig. 3a). Meanwhile, the distal end of Gα_s_ α5 helix moves 7 Å outward away from the TMD core relative to the equivalent Gα_q_ residue (measured at Cα atom of L^H5.25^, superscript refers to CGN system ^43^) (Fig. 3a). The G_q_ presents the largest solvent-accessible surface area (SASA, 1492 Å^2^) with the receptor compared to G_s_ (1293 Å^2^) and G_i_ (1167 Å^2^), consistent with a 6.6- to 20.3-fold increased potency of G_q_ coupling to CCK_A_R in comparison to that coupled with G_s_ and G_i_ (Extended Data Fig. Table 3). This finding supports the hypothesis that the size of the G protein coupling interface may correlate with the ability of a receptor to link with different G proteins ^23,24^. In addition, coupling of different G protein subtypes exhibits distinct effects on CCK-8 binding. Compared to G_s_ or G_i_ proteins, G_q_ coupling increases the binding affinity of CCK-8 (Extended Data Table 3), consistent with the increased binding activity of isoproterenol against β_2A_R in the presence of G_s_ protein ^16^. This finding indicates an allosteric modulation effect of G_q_ protein on CCK-8 binding, supporting the positive cooperativity between agonists and G proteins ^44^.

**Fig. 3.**
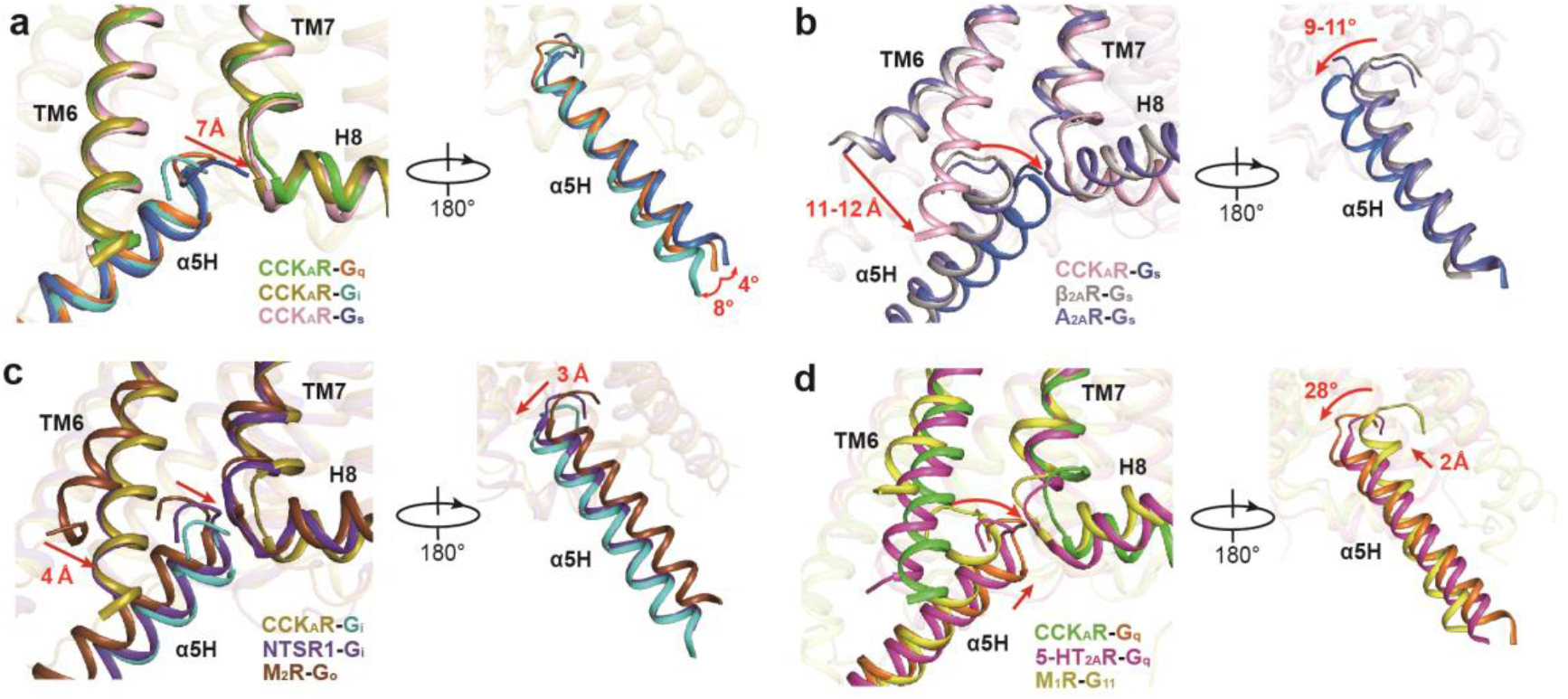
Structural comparison of TM6 and α5 helix between CCK_A_R-G protein complexes and representative G_s_-, G_q_-, and G_i_-coupled GPCR structures in two different views. **a**, Structural comparison of CCK_A_R–G_q_, CCK_A_R–G_s_, and CCK_A_R–G_i_ complexes. A 7 Å movement of the distal end of Gα_s_ α5 helix relative to that of Gα_q_ and swing of Gα α5 helix are highlighted as red arrows. **b**, Structural comparison of CCK_A_R–G_s_ with β_2A_R–G_s_ and A_2A_R–G_s_ complexes. Red arrows indicate an 11-12 Å displacement of TM6 and a 9-11° swing of Gα α5 helix of G_s_-coupled CCK_A_R relative to G_s_-coupled β_2A_R and A_2A_R. **c**, Structural comparison of CCK_A_R–G_i_ with NTSR1–G_i_ and M_2_R–G_o_ complexes. A 4 Å inward displacement of TM6 and a 3 Å Gα_i_ α5 helix shift of G_q_-coupled CCK_A_R in contrast to G_o_-coupled M_2_R are indicated as red arrows. **d**, Structural comparison of CCK_A_R–G_q_ with 5-HT_2A_R–G_q_ and M_1_R–G_11_ complexes. A 2 Å upward movement of Gα_q_ of G_q_-coupled CCK_A_R compared to G_q_-coupled 5-HT_2A_R and a 28° rotation relative to G_11_-coupled M_1_R are highlighted as red arrows. The complex structures are aligned based on TM2-TM4 of the receptors. β_2A_R–G_s_, A_2A_R–G_s_, NTSR1-G_i_, M_2_R–G_o_, 5-HT_2A_R–G_q_, and M_1_R–G_11_ structures (PDB codes: 3SN6, 5G53, 6OS9, 6OIK, 6WHA, and 6OIJ) are colored in gray, marine, purple blue, dark brown, magenta, and yellow, respectively.

In addition, comparisons of these three complex structures to previously reported G protein-coupled class A GPCRs reveal the different extent of TM6 displacement and concomitant shift of Gα α5 helix (Fig. 3b-d). TM6 of CCK_A_R in all three G protein complexes displays an 11-12 Å (measured at Cα atom of residue at position 6.27) smaller outward displacement in contrast to G_s_-coupled GPCRs, which translates into a notable swing of Gα α5 helix in the same direction (9-11° relative to G_s_-coupled β_2A_R and A_2A_R as measured at Cα atom of Y^H5.23^). This smaller displacement of TM6 is contrary to the previous assumption that TM6 of G_s_-coupled GPCRs undergoes a significant outward movement, thus opening a larger cytoplasmic pocket to accommodate bulkier residues at the distal end of Gα_s_ α5 helix relative to G_i/o_-coupled receptors ^23,45^. To avoid a potential clash with TM6, the distal end of the Gα_s_ α5 helix in the CCK_A_R–G_s_ complex stretches away from the TMD core and inserts into the crevice between TM6 and TM7–helix 8 joint. This featured conformation of Gα_s_ α5 helix in the CCK_A_R–G_s_ complex is unique compared to that in structures of the G_s_-coupled β_2A_R and A_2A_R, supporting the complexity of GPCR-G protein coupling mechanism (Fig. 3b).

TM6 and Gα α5 helix of CCK_A_R–G protein complexes display similar conformational changes to other G_i_- and G_q_-coupled GPCRs, such as the G_i_-coupled NTSR1 and the G_q_-coupled 5-HT_2A_R (Fig. 3c, d). TM6 of CCK_A_R–G_i_ protein complex is highly overlaid with that of G_i_-coupled NTSR1, while the cytoplasmic end of TM6 shows a 4 Å smaller outward displacement compared to that of G_o_-coupled M_2_R (Fig. 3c). On the G protein side, the α5 helix of Gα_i_ in the CCK_A_R–G_i_ complex shows a nearly overlapped conformation compared to that of the NTSR1–G_i_ complex. In contrast, it exhibits a 3 Å (measured at Cα atom of Y^H5.23^) shift away from TM6 relative to that of G_o_-coupled M_2_R (Fig. 3c). Structural comparison of G_q_-coupled CCK_A_R with G_q_/G_11_-coupled GPCRs demonstrates a 2 Å (measured at Cα atom of Y^H5.23^) upward toward the cytoplasmic cavity in contrast to the G_q_-coupled 5-HT_2A_R and a 28° rotation away from TM6 relative to G_11_-coupled M_1_R (Fig. 3d).

## Interaction patterns for the “wavy hook” of the CCK_A_R-G protein complexes

The “wavy hook” at the extreme C-terminus of the Gα α5 helix is thought to be one of the coupling specificity determinants for G protein ^46,47^, which undergoes distinct conformational rearrangements among the three CCK_A_R–G protein complexes (Fig. 3a).

A structural comparison of the interaction interface between the receptor cytoplasmic cavity and Gα “wavy hook” reveals distinct features of CCK_A_R–G protein coupling. Well-defined densities of Gα protein “wavy hook” residues allow for detailed structural analyses except for residues at the −1 position. L(−2)^H5.25^ in α5 helix is highly conserved across the G protein families and plays a pivotal role in G protein coupling. Both L358^H5.25^ in Gα_q_ and L353^H5.25^ in Gα_i_ hydrophobically interact with residues in TM3 and TM6 (R139^3.50^, I143^3.54^, V311^6.33^, and L315^6.37^) (Fig. 4a, b). Due to the notable displacement of Gα_s_ C-terminus, L393^H5.25^ in Gα_s_ moves 7 Å outward away from the TMD core relative to the equivalent Gα_q_ residue (Fig. 3a), repositioning it in a hydrophobic sub-pocket formed by M314^6.36^ and M373^7.56^ (Fig. 4c). In contrast to L(−2)^H5.25^, residues at positions H(−3)^5.24^, H(−4)^5.23^ and H(−5)^5.22^ are less conserved. N357(−3)^H5.24^ in Gα_q_ makes an H-bond with the backbone CO group of Y370^7.53^ (Fig. 4a). Owing to the replacement of Gα_i_ G352(−3)^H5.24^ and the reposition of Gα_s_ E392(−3)^H5.24^, the corresponding H-bond is absent in CCK_A_R–G_i_ and CCK_A_R–G_s_ complex structures. Additionally, Y356(−4)^H5.23^ in Gα_q_ forms extensive interactions with the receptor cytoplasmic cavity by making H-bonds with R139^3.50^ and Q153^ICL2^ (Fig. 4d). In contrast, C351(−4)^H5.23^ in Gα_i_ only forms a weak H-bond with R139^3.50^ via its backbone CO group (Fig. 4e). Y391(−4)^H5.23^ in Gα_s_ exhibits limited hydrophobic and Van der Waals interactions with residues in TM2 and TM3 (T76^2.39^, R139^3.50^, and A142^3.53^) (Fig. 4f). Furthermore, both E355(−5)^H5.22^ in Gα_q_ and D350(−5)^H5.22^ in Gα_i_ form salt bridges with R376^8.49^ in CCK_A_R, while Q390(−5)^H5.22^ in Gα_s_ disfavors the formation of corresponding electrostatic interaction (Fig. 4d-f). To understand the “wavy hook” mediated G-protein selectivity, we displaced the amino acids (H5.22-H5.25) in Gα_q_ subunit with the corresponding ones in Gα_s_ and Gα_i_ subunits. BRET assay results show that the Gα_i_ displacement has no impact on CCK_A_R-G protein coupling compared to wild-type Gα_q_ subunit. However, partially (E355Q or N357E) or completely Gα_s_ substitution remarkably decreased the G protein coupling activity of CCK_A_R (Fig. 4g). These results indicate that the “wavy hook” may play a crucial role in coupling selectivity of CCK_A_R with G_q_ over G_s_ protein.

**Fig. 4.**
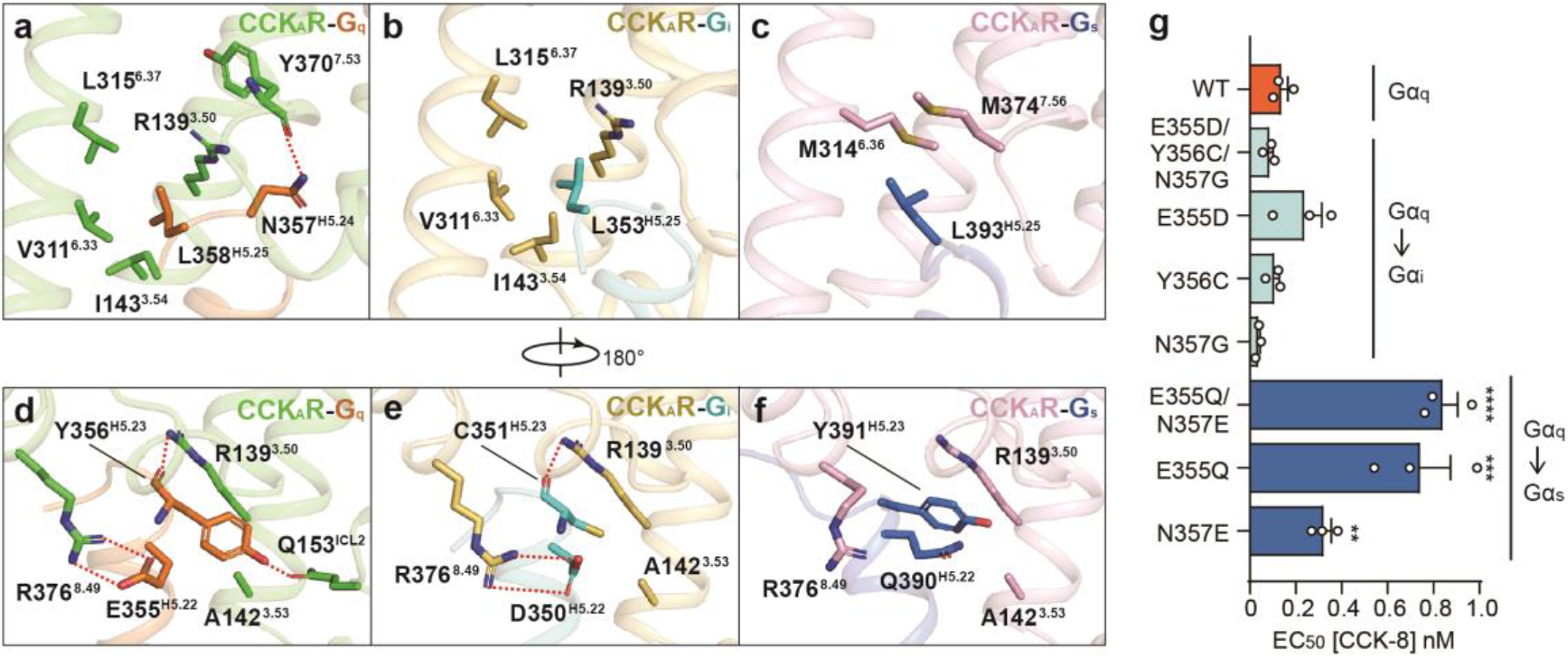
Distinct interaction patterns of residues from the “wavy hook” motif. **a-c,** Interaction details between CCK_A_R and L358^H5.25^ and N357^H5.24^ of Gα_q_ (**a**), L353^H5.25^ of Gα_i_ (**b**), and L393^H5.25^ of Gα_s_ subunit (**c**). **d-f**, Interaction details between CCK_A_R and Y356^H5.23^ and E355^H5.22^ of Gα_q_ (**d**), C351^H5.23^ and D350^H5.22^ of Gα_i_ (**e**), and Y391^H5.23^ and Q390^H5.22^ of Gα_s_ subunit (**f**). H-bonds and salt bridges are indicated as red dashed lines. **g**, BRET assay evaluating the effects of “wavy hook” substitutions on CCK_A_R-G protein coupling. The “wavy hook” residues of the Gα_q_ subunit were displaced by the corresponding residues in Gα_s_ and Gα_i_ subunits. All data were analyzed by one-way ANOVA. \*\*\*\**P*<0.0001, \*\*\**P*<0.001, \*\**P*<0.01.

## Contribution of CCK_A_R ICL3 to G_q_-coupling selectivity

In the CCK_A_R–G_q_ protein complex structure, CCK_A_R displays a comparable length of TM5 relative to M_1_R–G_11_ complex ^21^. However, the cytoplasmic end of CCK_A_R TM5 exhibits an 8 Å outward bend (measured at Cα atoms of A^5.73^), which prevents it from interacting with the Gα_q_ subunit (Fig. 5a). Instead, the ICL3 inserts into the cleft between TM5 of CCK_A_R and α5 helix of the Gα_q_ subunit (Fig. 5a). Compared to L225^5.75^ in M_1_R, I296^ICL3^ in CCK_A_R interacts with the same hydrophobic patch formed by side chains of Y325^S6.02^, F339^H5.06^, and A342^H5.09^ in Gα_q_ subunit, but is buried deeper to create more closely packed hydrophobic contacts (Fig. 5a, b). These hydrophobic interactions are critical to CCK_A_R–G_q_ coupling, as evidenced by our BRET analysis that I296^ICL3^G mutation significantly weakened G_q_ coupling to CCK_A_R but had no impact on G_s_ and G_i_ coupling (Fig. 5c, Extended Data Table 4). This hydrophobic patch that lies on the outer surface may be unique for the G_q/11_ subunit. The equivalent residues in Gα_s_ and Gα_i_ subunits are polar or charged residues, which would be energetically unfavorable to form hydrophobic interactions (Extended Data Fig. 10). Indeed, this unconventional ICL3-G_q_ interaction is not seen in the structures of G_s_- and G_i/o_-coupled CCK_A_Rs (Fig. 3b, c). Together, our findings offer structural evidence on the possible role of ICL3 in CCK_A_R–G_q_ coupling preference. Hydrophobic residues on the inner surface of the ICL3 loop of CCK_A_R or the extended TM5 of M_1_R may represent a common feature of G_q/11_-coupled GPCRs.

**Fig. 5.**
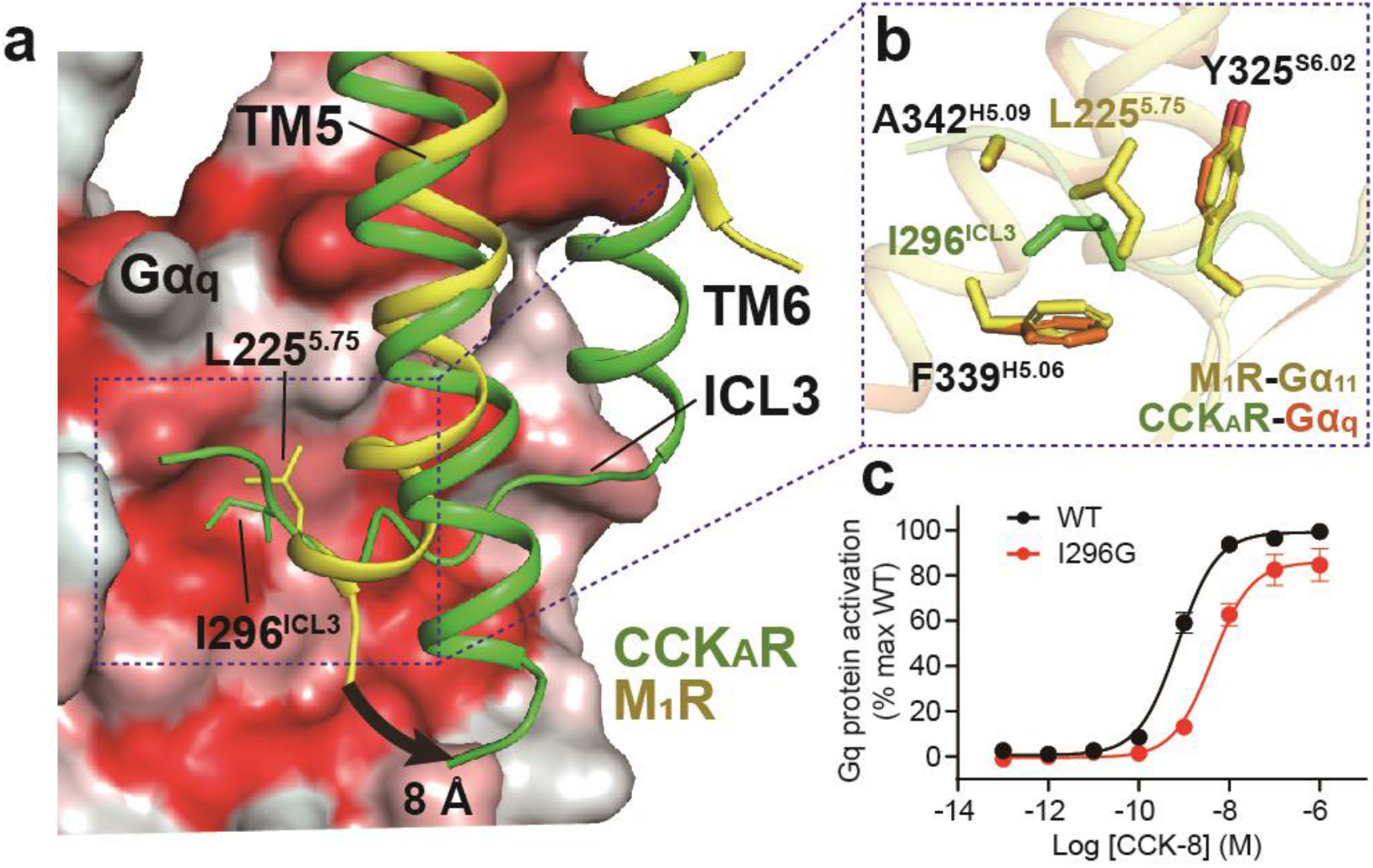
Interaction between ICL3 loop of CCK_A_R and Gα_q_ subunit. **a,** I296^ICL3^ of CCK_A_R and L225^5.75^ of M_1_R occupy the same hydrophobic sub-pocket of the Gα subunit. The Gα_q_ subunit is shown as a surface presentation by hydrophobicity (hydrophobic surface in red). An 8 Å outward bend of TM5 of CCK_A_R relative to that of M_1_R is highlighted by a black arrow. **b**, Detailed interactions between I296^ICL3^(CCK_A_R)/L225^5.75^(M_1_R) and hydrophobic patch comprised by Y325^S6.02^, F339^H5.06^, and A342^H5.09^ of Gα_q_ and Gα_11_ subunits. **c**, BRET assay indicates that I296G mutation decreases the association rate of CCK_A_R with G_q_ heterotrimer (n=3). WT, wild-type.

## Conclusions

As the largest family of cell surface receptors, GPCRs have more than 800 members but only couple to four G protein subtypes. Specific GPCR signaling requires the receptor to couple with either a single or multiple G protein subtypes ^47–49^. Thus, one of the main questions is how does a given GPCR select a G protein subtype for downstream signal transduction. The critical G protein determinants of selectivity vary widely for different receptors that couple to specific G proteins. It is thought that G_s_- or G_q_-coupled receptors are relatively promiscuous and to some extent couple to G_i1_ ^24^. However, G_i_-coupled receptors are more selective ^24^. The minor outward movement of TM6 contributes to such a superior G_i_-coupling selection as opposed to that of G_s_ ^18,25,46,50,51^. Although proven to be promiscuous, G_q_-coupled receptors tend to adopt an active conformation similar to that of G_i_-coupled GPCRs, reflecting the complexity of the GPCR-G protein coupling mechanism ^21,22^. Since CCK_A_R has the ability to couple with different G protein subtypes, it stands out as a suitable model for studying the promiscuity of G protein coupling. In this paper, we show that TM6 of CCK_A_R undergoes a similar outward displacement relative to G_i/o_-coupled (NTSR1 and M_2_R) and G_q/11_-coupled GPCRs (5-HT_2A_R and M_1_R) but has a smaller shift relative to G_s_-coupled GPCRs (β_2A_R and A_2A_R). CCK_A_Rs share almost identical conformations, whereas G_q_, G_s_, and G_i_ proteins vary in distinct orientations, producing different sizes of receptor-G protein interface. The predominant coupling to G_q_ by CCK_A_R can be explained by the largest interface among three CCK_A_R-G protein complexes. Structural comparison of the three CCK_A_R–G protein complexes reveals that “wavy hook” residues of Gα α5 helix and ICL3 of the receptor are important for the coupling promiscuity. In addition, detailed inspections disclose structural clues relative to the recognition mechanism of sulfated CCK-8 by CCK_A_R, in which R197^ECL2^ is a major determinant. Together, our structures provide a framework for better understanding of ligand recognition as well as G protein coupling selectivity and promiscuity by CCK_A_R.

## Methods

### Expression and purification of CCK_A_R-G protein complexes

The WT CCK_A_R (residues 1-428) was applied for cryo-EM studies. The full-length CCK_A_R cDNA was cloned into a modified pFastBac vector (Invitrogen) containing a hemagglutinin (HA) signal sequence followed by an 8× histidine tag, a double-MBP tag, and a TEV protease site before the receptor sequence using homologous recombination (CloneExpress One Step Cloning Kit, Vazyme) (Extended Data Fig. 2a). The N-terminal 1-29 amino acids of Gα_q_ was replaced by the equivalent residues of Gα_i1_ to facilitate the scFv16 binding ^21^. An engineered Gα_s_ construct was generated based on mini-Gα_s_ ^33^. The N-terminal 1-18 amino acids and α-helical domain of Gα_s_ were replaced by human Gα_i1_, thus providing binding sites for scFv16 and Fab-G50, respectively ^18,21^. Additionally, human Gα_i1_ with two dominant-negative mutations (G203A and A326S ^34^) was used to assemble a stable GPCR-G_i_ protein complex. These two cognate mutations also exist in engineered Gα_q_ and Gα_s_ (Extended Data Fig. 2b). Receptor, rat H6-Gβ, bovine Gγ, and the specific Gα subunit were co-expressed in *Spodoptera frugiperda* (*sf9*) insect cells (Invitrogen) as previously described ^52^. In addition, GST-Ric-8A (a gift from Dr. B. Kobilka) was applied to improve the expression of Gα_q_.

ScFv16 was applied to improve the protein stability of CCK_A_R–G_q_ and CCK_A_R–G complex samples. The monomeric scFv16 was prepared as previously reported ^53^. Cell pellets of the co-expression culture were thawed and lysed in 20 mM HEPES, pH 7.4, 100 mM NaCl, 10% glycerol, 5 mM MgCl_2_, and 10 mM CaCl_2_ supplemented with EDTA-free protease inhibitor cocktail (TargetMol). CCK_A_R–G protein complexes were assembled at room temperature (RT) for 1 h by the addition of 10 μM CCK-8 (GenScript) and 25 mU/mL apyrase. Then the lysate was solubilized in 0.5% LMNG, 0.1% CHS, and the soluble fraction was purified by nickel affinity chromatography (Ni Smart Beads 6FF, SMART Lifesciences). In the case of CCK_A_R–G_i_ and CCK_A_R–G_q_ complexes, a 3 molar excess of scFv16 was added to the protein elute. The mixture was incubated with amylose resin for 2 h at 4°C. The excess G protein and scFv16 were washed with 20 column volumes of 20 mM HEPES, pH 7.4, 100 mM NaCl, 10% glycerol, 0.01% LMNG, 0.002% CHS, and 2 μM CCK-8. TEV protease was then included to remove the N terminal fusion tags of CCK_A_R. After 1 h incubation at RT, the flow-through was collected, concentrated, and injected onto a Superdex 200 10/300 column equilibrated in the buffer containing 20 mM HEPES, pH 7.4, 100 mM NaCl, 0.00075% LMNG, 0.00025% GDN, 0.0002% CHS, and 10 μM CCK-8. The monomeric complex peak was collected and concentrated to about 5 mg/mL for cryo-EM studies.

### Cryo-EM grid preparation and image collection

For preparation of cryo-EM grids, 2.5 μL of each purified CCK_A_R–G protein complex was applied individually onto the glow-discharged holey carbon grids (Quantifoil, Au300 R1.2/1.3) in a Vitrobot chamber (FEI Vitrobot Mark IV). The Vitrobot chamber was set to 100% humidity at 4°C. Extra samples were blotted for 2 s and were vitrified by plunging into liquid ethane. Grids were stored in liquid nitrogen for condition screening and data collection usage.

Automatic data collection of CCK-8–CCK_A_R–G protein complexes were performed on a FEI Titan Krios operated at 300 kV. The microscope was operated with a nominal magnification of 81,000× in counting mode, corresponding to a pixel size of 1.045 Å for the micrographs. A total of 5,415 movies for the dataset of CCK-8–CCK_A_R–G_q_-scFv16 complex, 5008 movies for the dataset of CCK-8–CCK_A_R–G_s_ complex, and 4,811 movies for the dataset of CCK-8–CCK_A_R–G_i_-scFv16 complex were collected, respectively, by a Gatan K3 Summit direct electron detector with a Gatan energy filter (operated with a slit width of 20 eV) (GIF) using the SerialEM software. The images were recorded at a dose rate of about 26.7 e/Å^2^/s with a defocus ranging from −0.5 to −3.0 μm. The total exposure time was 3 s and intermediate frames were recorded in 0.083 s intervals, resulting in a total of 36 frames per micrograph.

### Image processing and map reconstruction

Image stacks were subjected to beam-induced motion correction and aligned using MotionCor 2.1. Contrast transfer function (CTF) parameters were estimated by Ctffind4. The data processing was performed using RELION-3.0 ^54^. The micrographs with the measured resolution worse than 4.0 Å and micrographs imaged within carbon area were discarded, generating 3,806 micrographs for CCK-8–CCK_A_R–G_q_–scFv16 dataset, 4,963 micrographs for CCK-8–CCK_A_R–G_s_ dataset, and 4,543 micrographs for CCK-8–CCK_A_R–G_i_–scFv16 dataset for further data processing. Particle selection, 2D and 3D classifications were performed on a binned dataset with a pixel size of 2.09 Å. About 2,000 particles were manually selected and subjected to 2D classification. Representative averages were chosen as template for particle auto-picking. The auto-picking process produced 3,405,355 particles for CCK-8–CCK_A_R–G_q_–scFv16 complex, 4,680,972 particles for CCK-8–CCK_A_R–G_s_ complex, and 4,270,010 particles for CCK-8–CCK_A_R–G_i_–scFv16 complex, which were subjected to reference-free 2D classifications to discard bad particles. Initial reference map models for 3D classification were generated by Relion using the representative 2D averages. For CCK-8–CCK_A_R–G_q_–scFv16 complex, the particles selected from 2D classification were subjected to 6 rounds 3D classifications, resulting in a single well-defined subset with 555,628 particles. For CCK-8–CCK_A_R–G_s_ complex, the particles resulting from 2D classification were subjected to 5 rounds 3D classifications, resulting in two well-defined subsets with 499,924 particles. For CCK-8–CCK_A_R–G_i_-scFv16 complex, the particles selected from 2D classification were subjected to 7 rounds 3D classifications, resulting in two well-defined subsets with 140,602 particles. Further 3D refinement, CTF refinement, Bayesian polishing and DeepEnhancer processing generated density maps with an indicated global resolution of 2.9 Å for CCK-8–CCK_A_R–G_q_–scFv16 complex, 3.1 Å for CCK-8–CCK_A_R–G_s_ complex, and 3.2 Å for CCK-8–CCK_A_R–G_i_–scFv16 complex, respectively, at a Fourier shell correlation of 0.143.

### Model building and refinement

For the CCK_A_R–G_q_ complex, the initial G_q_ protein and scFv16 model were adopted from the cryo-EM structure of the M_1_R–G_11_ protein complex (PDB: 6OIJ) ^21^. The initial CCK_A_R model was generated by an online homology model building tool ^55^. All models were docked into the EM density map using Chimera ^56^, followed by iterative manual adjustment and rebuilding in COOT ^57^ and ISOLDE ^58^, and real-space refinement using Phenix programs ^59^. The model statistics were validated using Phenix comprehensive validation. A model of the refined CCK_A_R from the CCK_A_R–G_q_ complex was used for the other two complexes. Models from PTH1R–G_s_ (PDB: 6NBF) and FPR2–G_i_ (PDB: 6OMM) were used as templates for the model building of G_s_ in the CCK_A_R–G_s_ complex and G_i1_–scFv16 in the CCK_A_R–G_i_ complex, respectively. Then the fitted models were built the same way as the CCK_A_R–G_q_ complex. The final refinement statistics are provided in Extended Data Table 1.

### Radiolabeled ligand-binding assay

The WT or mutant CCK_A_Rs were transiently transfected into HEK 293T/17 cells (purchased from the Cell Bank at the Chinese Academy of Sciences) which were cultured in poly-D-lysine coated 96-well plate. Twenty-four h later, the cells were washed twice and incubated with blocking buffer (DMEM medium supplemented with 33 mM HEPES, and 0.1% (w/v) BSA, pH 7.4) for 2 h at 37°C. After three times washes by cold-ice PBS, the cells were treated by a constant concentration of ^125^I-CCK-8 (40 pM, PerkinElmer) plus 8 different doses of CCK-8 (1 pM to 10 μM) for 3 h at RT. Cells were washed three times with ice-cold PBS and lysed by 50 μL lysis buffer (PBS supplemented with 20 mM Tris-HCl and 1% (v/v) Triton X-100, pH 7.4). Subsequently, the plates were counted for radioactivity (counts per minute, CPM) in a scintillation counter (MicroBeta^2^ plate counter, PerkinElmer) using 150 μl scintillation cocktail (OptiPhase SuperMix, PerkinElmer).

### G protein dissociation assay

G protein dissociation was monitored by BRET (bioluminescence resonance energy transfer) experiments performed as previously reported ^60^. Briefly, a C-terminal fragment of the GRK3 (GRK3ct) fused to a luciferase serves as a BRET donor. Gβγ dimer is labeled with a fluorescent protein Venus, a BRET acceptor. Upon G protein heterotrimer activation, free Gβγ-Venus is released and binds to membrane-associated GRK3ct-luciferase, leading to an increased signal detectable by BRET.

HEK 293T/17 cells were seeded onto 10 μg/mL Matrigel-coated 6-well plate (1×10^6^ cells/well). After 4 h culture, WT or mutant CCK_A_R (0.84 μg), Gα (Gα_q_, Gα_s_, and Gα_i_, 2.1 μg each), Gβ (0.42 μg), Gγ (0.42 μg), and GRK (0.42 μg) were transiently transfected with Lipofectamine™ LTX Reagent (Invitrogen). Twenty-four hours post-transfection, cells were washed once with DMEM medium (no phenol red) and detached by EDTA. Cells were then harvested with centrifugation at 1000 rpm for 5 min and resuspended in DMEM medium. Approximately 75,000 cells per well were distributed in 96-well flat-bottomed white microplates (PerkinElmer). The NanoBRET substrate (furimazine, 25 μL/well, Promega) was added, and the BRET signal (535 nm/475 nm ratio) was determined using an EnVision multilabel plate reader (PerkinElmer). The average baseline value recorded before CCK-8 stimulation was subtracted from BRET signal values.

### NanoBiT G-protein recruitment assay

The recruitment of CCK_A_R to G_i_-protein was detected in *sf9* cells using NanoBiT method as previously reported ^61^. Briefly, the LgBiT fragment of NanoBiT luciferase was fused to the C-terminus of CCK_A_R. SmBiT was fused to the C-terminus of Gβ subunit with a 15-amino acid flexible linker. CCK_A_R-LgBiT, Gα_i1_, SmBiT-fused human Gβ1 and human Gγ2 were co-expressed in *sf9* insect cells. Cell pellets were collected by centrifugation after infection for 48 h. The cell suspension was dispensed in a 96-well plate (64,000 cells per well) at a volume of 80 μL diluted in the assay buffer (HBSS buffer supplemented with 10 mM HEPES, pH 7.4) and incubated for 30 min at 37°C. The cells were then reacted with 10 μL of 50 mM coelenterazine H (Yeasen) for 2h at RT. Luminescence signal was measured using an EnVision plate reader (PerkinElmer) at 30 s intervals (25°C). The baseline was measured before CCK-8 addition for 8 intervals, and the measurements continued for 20 intervals following ligand addition. Data were corrected to baseline measurements and the results were analyzed using GraphPad Prism 8.0 (Graphpad Software Inc.).

### NanoBiT G-protein dissociation assay

G_s_ activation was measured by a NanoBiT dissociation assay. G protein NanoBiT split luciferase constructs were generated by fusing the LgBiT in Gα_s_ and the SmBiT to Gγ (a gift from Dr. Asuka Inoue, Tohoku University) as previously reported ^62^. In brief, HEK 293T/17 cells were plated in 10 cm plates at a density of 3×10^6^ cells per plate. After 24 h, cells were transfected with 1.62 μg plasmids of receptor, 0.81 μg Gα_s_-LgBiT, 4.1 μg Gβ, and 4.1 μg SmBiT-Gγ using Lipofectamine™ LTX Reagent (Invitrogen). The transiently transfected cells were then seeded into poly-D-lysine coated 96-well plates (50,000 cells per well) and grown overnight before incubation in an assay buffer. The measurement of luminescence signal was identical to the steps described above.

### Surface expression assay

HEK 293T/17 cells were seeded into a 6-well plate and incubated overnight. After transient transfection with WT or mutant plasmids for 24 h, the cells were collected and blocked with 5% BSA in PBS at RT for 15 min and incubated with primary anti-Flag antibody (1:300, Sigma-Aldrich) at RT for 1 h. The cells were then washed three times with PBS containing 1% BSA followed by 1 h incubation with donkey anti-mouse Alexa Fluor 488-conjugated secondary antibody (1:1000, ThermoFisher) at 4°C in the dark. After three washes, the cells were resuspended in 200 μl of PBS containing 1% BSA for detection in a NovoCyte flow cytometer (ACEA Biosciences) utilizing laser excitation and emission wavelengths of 488 nm and 519 nm, respectively. For each assay point, approximately 15,000 cellular events were collected, and the total fluorescence intensity of positive expression cell population was calculated.

### Molecular docking

Before docking, hydrogens were added to CCK_A_R and the whole system coordinates were optimized with a pH of 7.0. A grid file was then generated on the peptide pocket in our G_q_-coupled CCK_A_R structure. Small molecule ligands Glaxo-11p, GW-5823, and CE-326597 were prepared in the OPLS3 force field with a pH of 7.0 to generate 3D structures. Finally, glide docking with standard precision was applied to all ligands and the structures with the best docking score were picked as outputs.

## Acknowledgments

We thank Kirill A. Martemyanov for expert advice on BRET assay. The cryo-EM data were collected at the Cryo-Electron Microscopy Research Center, Shanghai Institute of Materia Medica (SIMM). The authors thank the staff at the SIMM Cryo-Electron Microscopy Research Center for their technical support.

## Author contributions

Q.F.L. screened the expression constructs, optimized the CCK_A_R-G protein complexes, prepared the protein samples for final structure determination, participated in cryo-EM grid inspection and data collection, built and refined the structure models, prepared the constructs for functional assays, analyzed the structures, and prepared the figures and wrote the initial manuscript; Y.W.Z. performed cryo-EM grid preparation, data collection, structure determination, and participated in protein sample optimization, figure and manuscript preparation; T.C. helped build and refine the structure model; X.H.H. performed the molecular docking; J.D. and W.C.Y. designed G protein constructs; F.L.Z. participated in data analysis; B.L.W. and Q.Z. participated in research supervising; H.E.X. conceived and supervised the project, analyzed the structures, and initiated collaborations with M.-W.W., supervised Q.F.L., Z.Y.W., F.L.Z., J.D., W.C.Y.; M.-W.W and D.H.Y. supervised X.Q.C., A.T.D., and C.Y.Y. in G protein assay development and data analysis; M.-W.W. participated in manuscript writing; Y.J. supervised the studies, performed the structural analysis, and prepared the figures and wrote the manuscript with input from all co-authors.

## Data availability

All data is available in the main text or the supplementary materials. Materials are available from the corresponding authors upon reasonable request.

---

**Extended Data Fig. 1.**
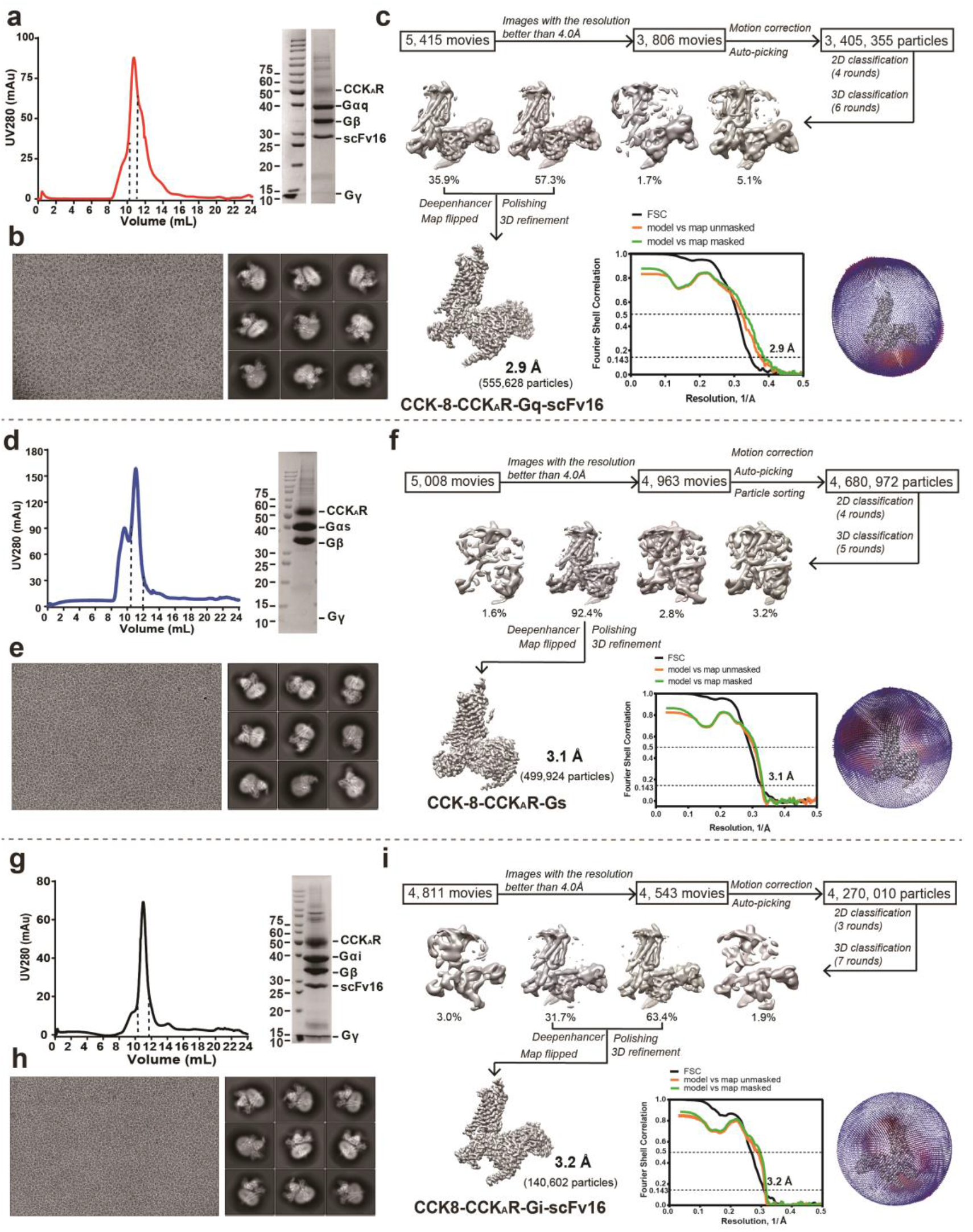
Cryo-EM workflows for structure determination of CCK_A_R–G protein complexes. **a,** Size exclusion chromatography (SEC) profile and SDS-PAGE analysis of the CCK-8–CCK_A_R–G_q_–scFv16 protein complex sample. **b**, Representative cryo-EM micrograph and 2D classification averages of the CCK-8–CCK_A_R–G_q_–scFv16 complex, the 2D averages display different secondary features in different views. **c**, Single-particle cryo-EM data processing flowcharts of the CCK-8–CCK_A_R–G_q_–scFv16 by Relion 3.1, including the Euler angle distribution of particles used in the final refinement and the fourier shell correlation (FSC) curves. The global resolution defined at the FSC=0.143 is 2.9 Å. **d**, Size exclusion chromatography (SEC) profile and SDS-PAGE analysis of the CCK-8–CCK_A_R–G_s_ protein complex sample. **e**, Representative cryo-EM micrograph and 2D classification averages of the CCK-8–CCK_A_R–G_s_ complex. **f**, Single-particle cryo-EM data processing flowcharts of the CCK-8–CCK_A_R–G_s_ by Relion 3.0, including the Euler angle distribution of particles used in the final refinement and the fourier shell correlation (FSC) curves. The global resolution defined at the FSC=0.143 is 3.1 Å. **g**, Size exclusion chromatography (SEC) profile and SDS-PAGE analysis of the CCK-8–CCK_A_R–G_i_–scFv16 protein complex sample. **h**, Representative cryo-EM micrograph and 2D classification averages of the CCK-8–CCK_A_R–G_i_–scFv16 complex. **i**, Single particle cryo-EM data processing flowcharts of the CCK-8–CCK_A_R–G_i_–scFv16 by Relion 3.0, including the Euler angle distribution of particles used in the final refinement and the fourier shell correlation (FSC) curves. The global resolution defined at the FSC=0.143 is 3.2 Å.

**Extended Data Fig. 2.**
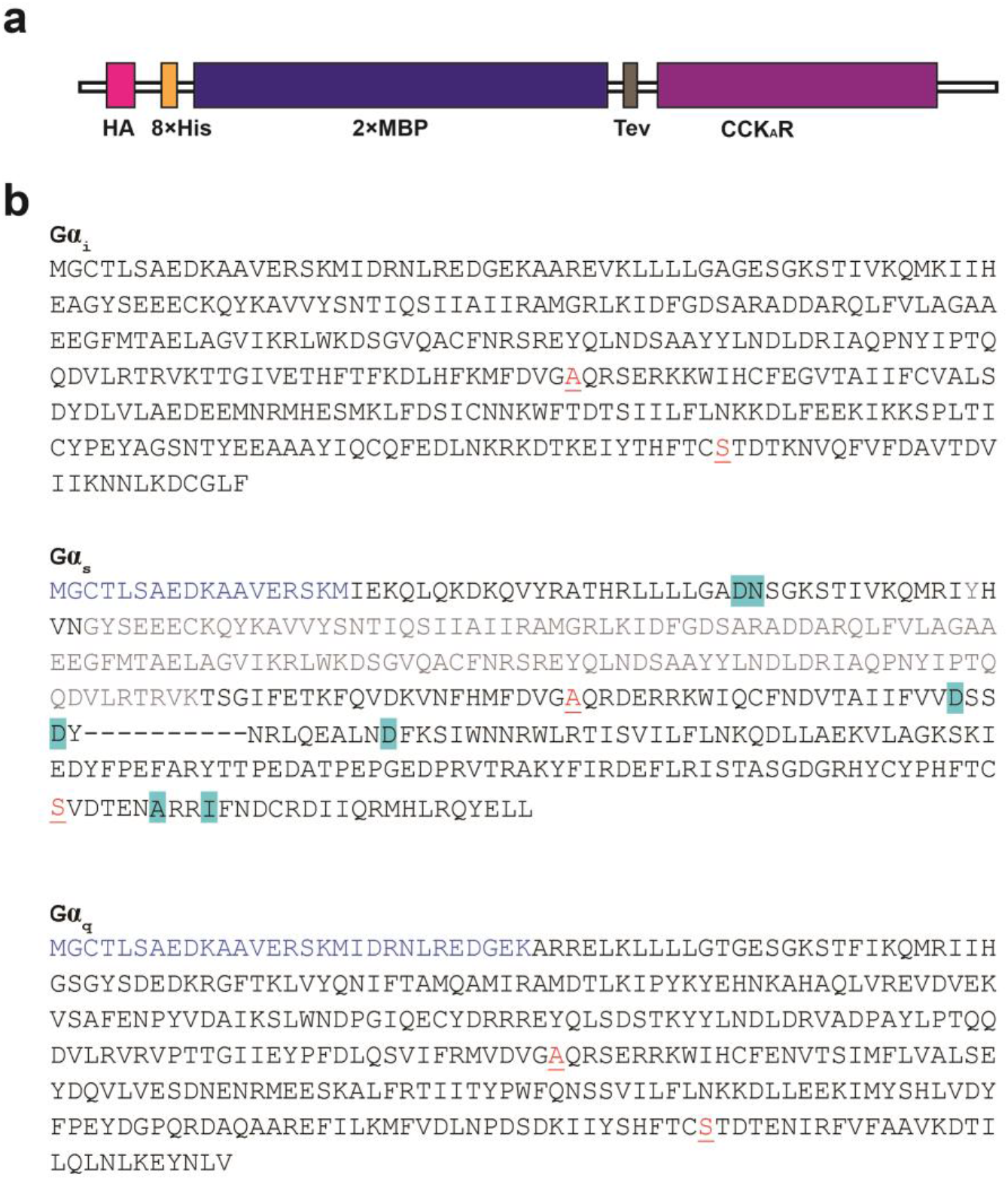
Receptor and Gα subunits used in the cryo-EM structure determination. **a**, A schematic illustration of the CCK_A_R construct used in cryo-EM studies. HA, hemagglutinin signal sequence; 2×MBP, double-MBP tag. **b**, Protein sequences of Gα_q_, Gα_s_, and Gα_i1_ subunits. N-terminal sequence replaced in Gα_s_ and Gα_q_ is shown in blue. The two dominant-negative mutations are colored red and underlined. Stabilization mutations derived from the reported mini-Gα_s_ are highlighted in cyan. AHD domain of the Gα_s_ is replaced with the equivalent region of Gα_i1_ and colored in gray.

**Extended Data Fig. 3.**
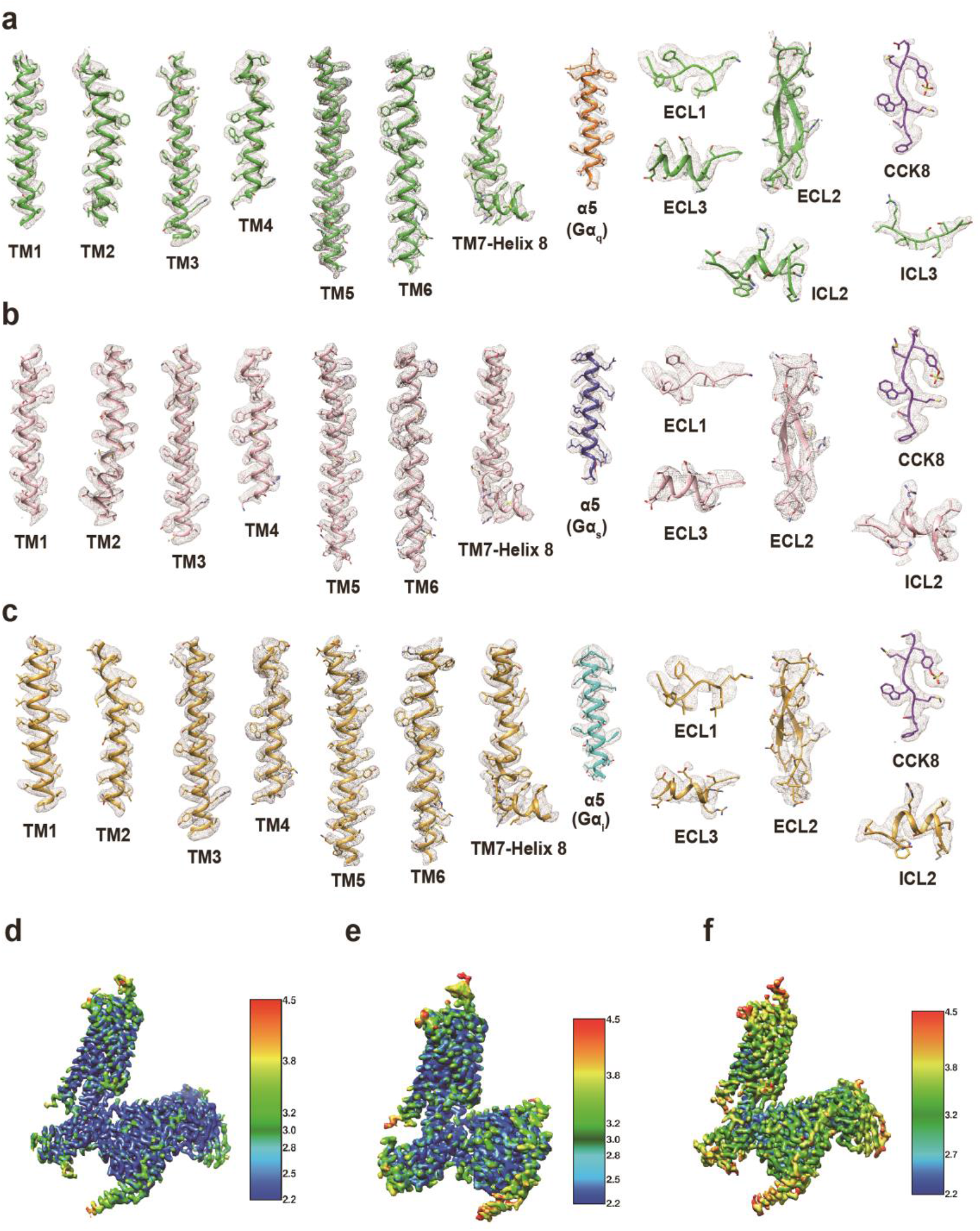
Local cryo-EM density maps of CCK_A_R–G protein complexes. **a,** Cryo-EM density maps of TM1-TM7, ECL1-ECL3, ICL2, ICL3, CCK-8 peptide and α5 helix of Gα_q_ in the CCK-8–CCK_A_R–G_q_–scFv16 structure. **b**, Cryo-EM density maps of TM1-TM7, ECL1-ECL3, ICL2, CCK-8 peptide and α5 helix of Gα_s_ in the CCK-8–CCK_A_R–G_s_ structure. **c**, Cryo-EM density maps of TM1-TM7, ECL1-ECL3, ICL2, CCK-8 peptide and α5 helix of Gα_i_ in the CCK-8–CCK_A_R–G_i_–scFv16 structure. **d-f**, The global density maps of the CCK-8–CCK_A_R–G_q_–scFv16 (**d**), CCK-8–CCK_A_R–G_s_ (**e**), and CCK-8–CCK_A_R–G_i_–scFv16 (**f**) colored by local resolution (Å). The density maps are shown at thresholds of 0.08, 0.055 and 0.05 for the CCK_A_R–G_q_, CCK_A_R–G_s_ and CCK_A_R–G_i_ complex, respectively.

**Extended Data Fig. 4.**
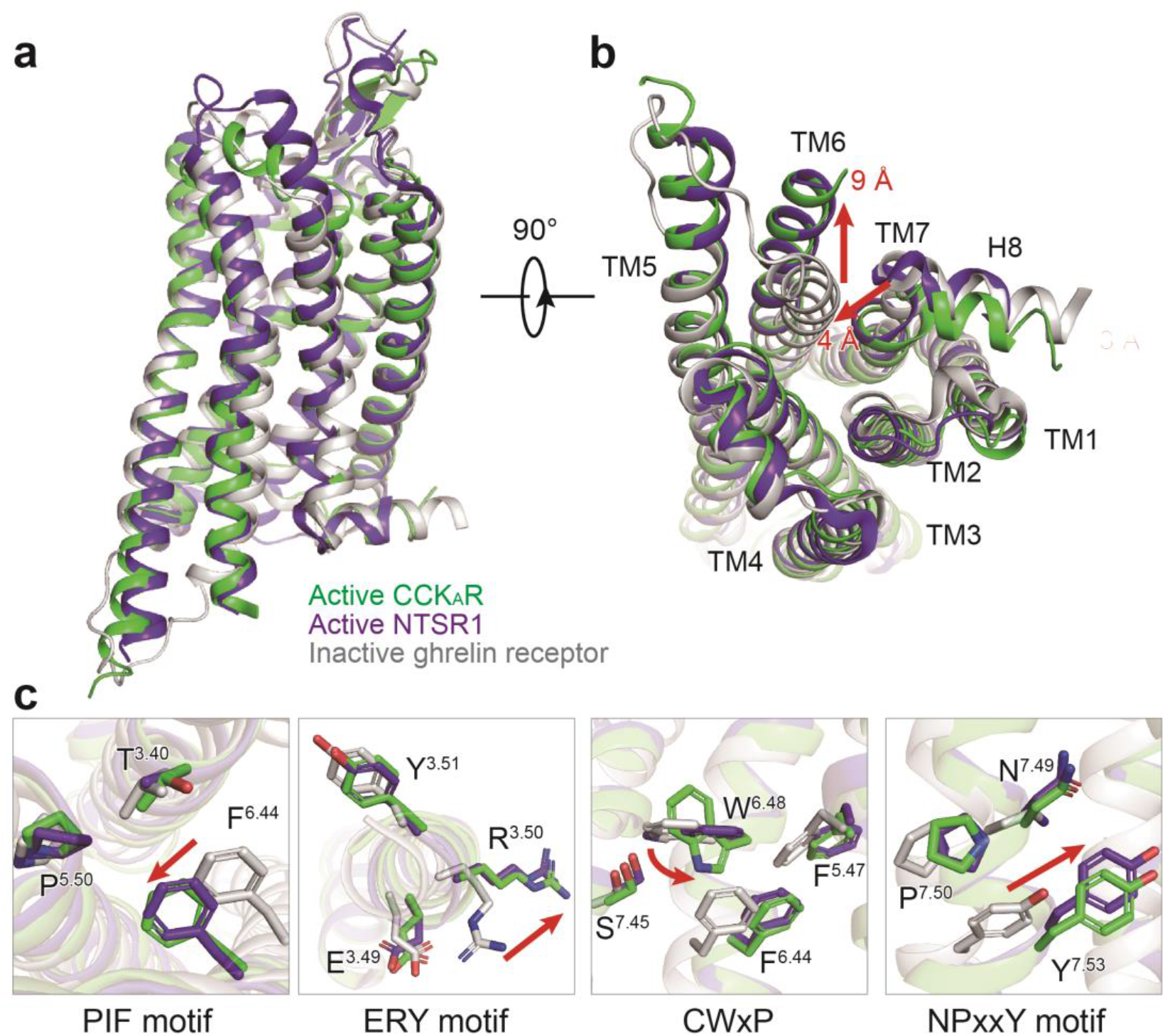
Active conformation of CCK_A_R. **a-b,** Structural comparison of inactive ghrelin receptor (grey), active NTSR1 (purple blue), and active CCK_A_R (green). Side view (**a**) and intracellular view (**b**) of the overall comparison are shown. **c,** Structural rearrangements of key activation motifs (PIF, ERY, CWxP, and NPxxY) in CCK_A_R compared to inactive GHSR and active NTSR1. NTSR1, neurotensin receptor 1.

**Extended Data Fig. 5.**
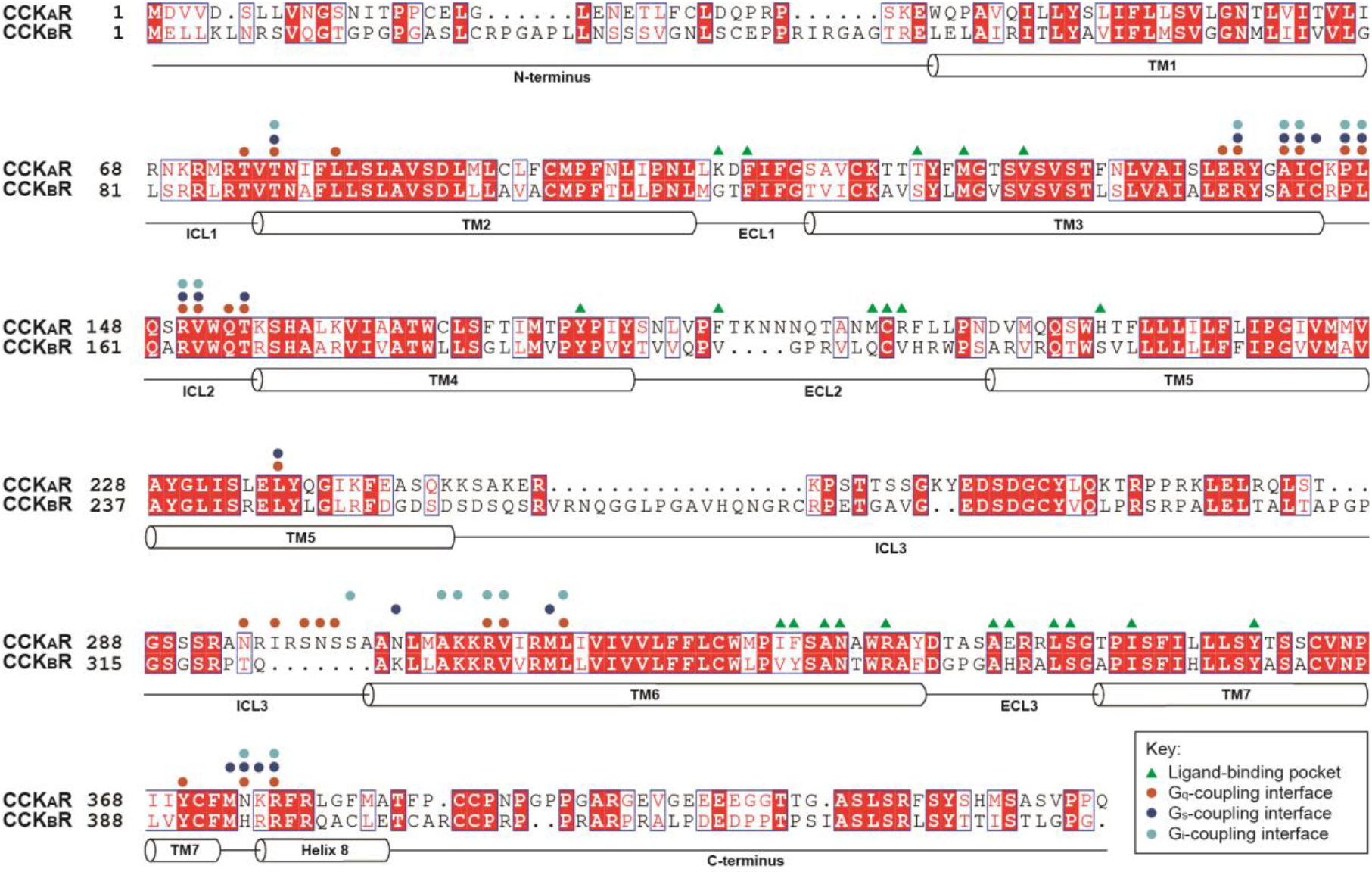
Sequence alignment of CCK receptors. Helical secondary structures are shown based on CCK_A_R. Residues involved in ligand-binding are labeled with green triangles. Residues involved in G protein coupling are labeled with circles (orange, G_q_; blue, G_s_; cyan, G_i_).

**Extended Data Fig. 6.**
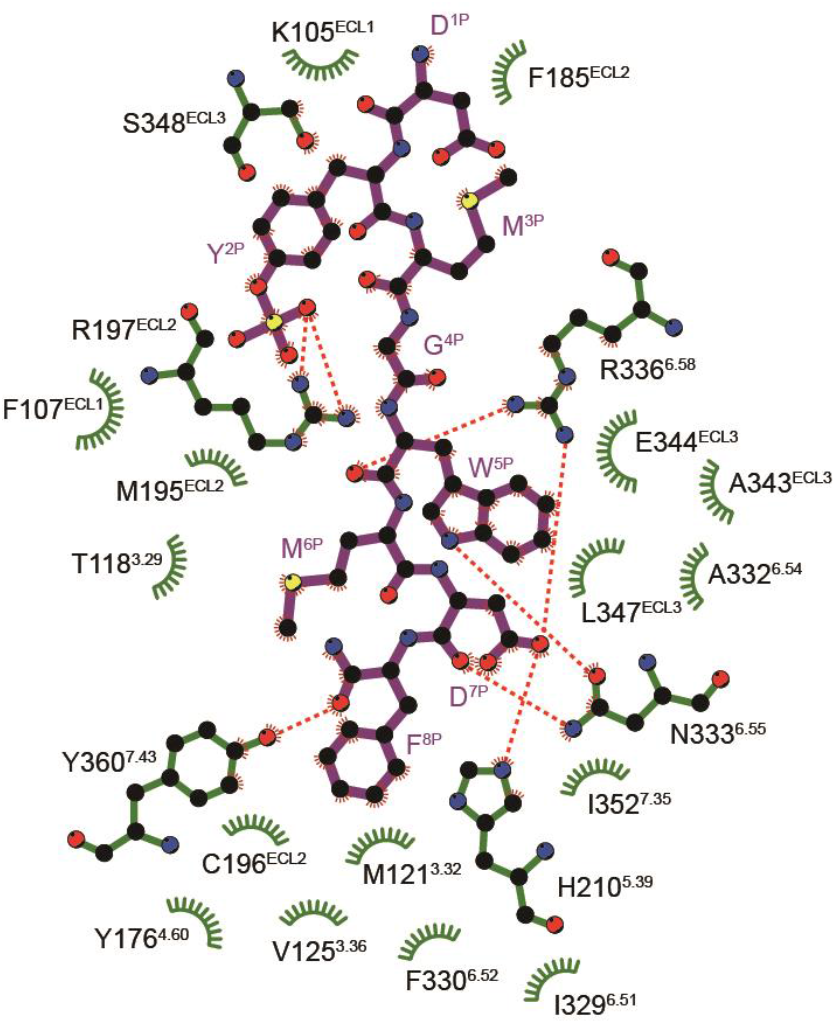
2D interaction plot of CCK_A_R recognition by sulfated CCK-8. Residues in the ligand-binding pocket are colored in green. CCK-8 is displayed as magenta sticks. Polar interactions are indicated as red dashed lines.

**Extended Data Fig. 7.**
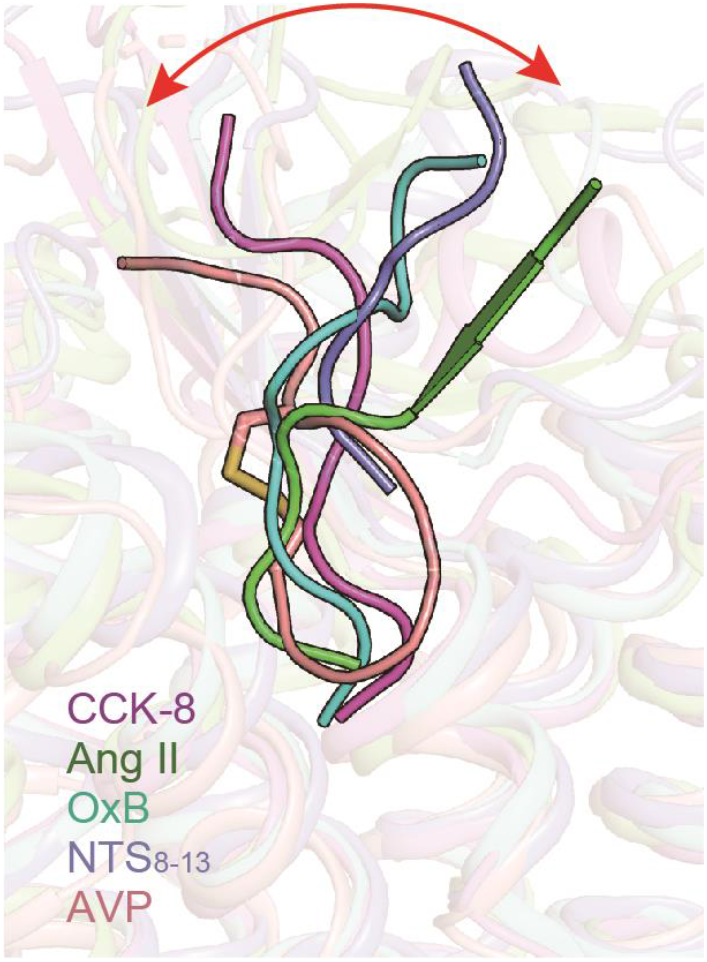
Structure comparison of CCK-8 with other neuropeptides solved to date. The neuropeptides are shown as a cartoon. The shift of the extracellular part of neuropeptides is highlighted as a red arrow. CCK-8 in the CCK-8–CCK_A_R–G_q_ complex structure, magenta; Ang II, angiotensin II (PDB: 6OS0), green; OxB, orexin B (PDB: 7L1U), cyan; NTS_8-13_, neurotensin 8-13 (PDB: 6OS9), purple blue; AVP, arginine vasopressin (PDB: 7DW9), salmon.

**Extended Data Fig. 8.**
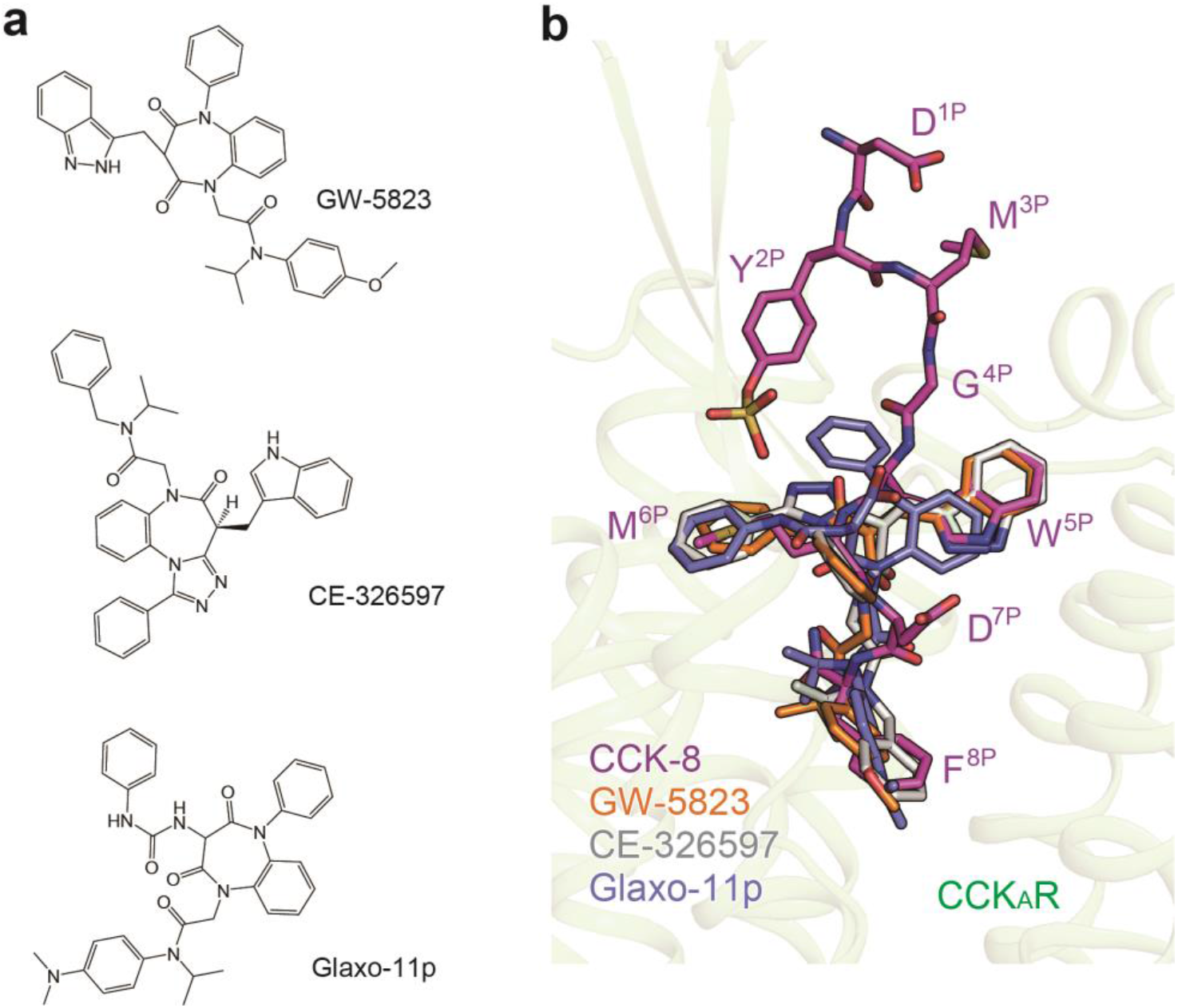
Molecular docking of small molecule agonists to the CCK_A_R structure. **a,** Chemical structures of small molecule agonists of CCK_A_R. **b,** Comparison of the binding poses of three agonists with CCK-8. CCK-8, magenta; GW-5823, orange; CE-326597, grey; Glaxo-11p, purple blue. CCK-8 and small molecule agonists are shown as sticks. The amino acids of CCK-8 are labelled.

**Extended Data Fig. 9.**
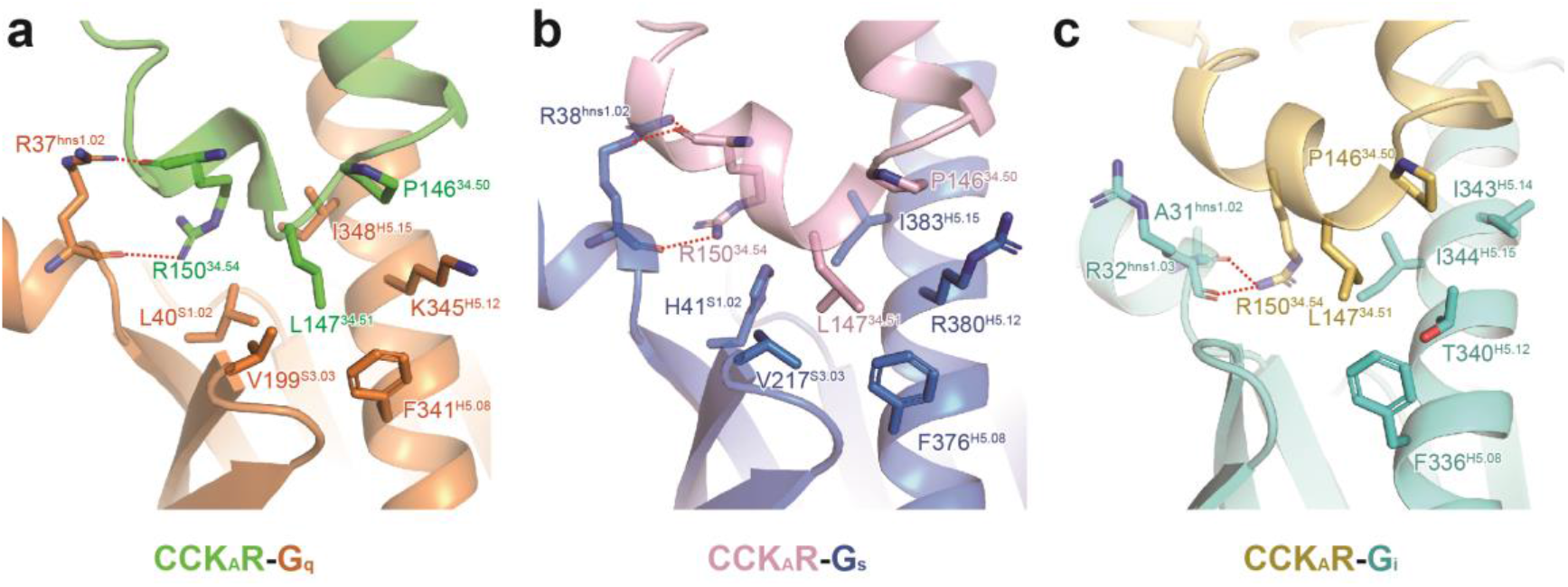
The interface between CCK_A_R ICL2 and different G proteins. Detailed interaction between the receptor and Gα_q_ (**a**), Gα_s_ (**b**), and Gα_i_ (**c**) are shown. Side chains of related residues are shown as sticks.

**Extended Data Fig. 10.**
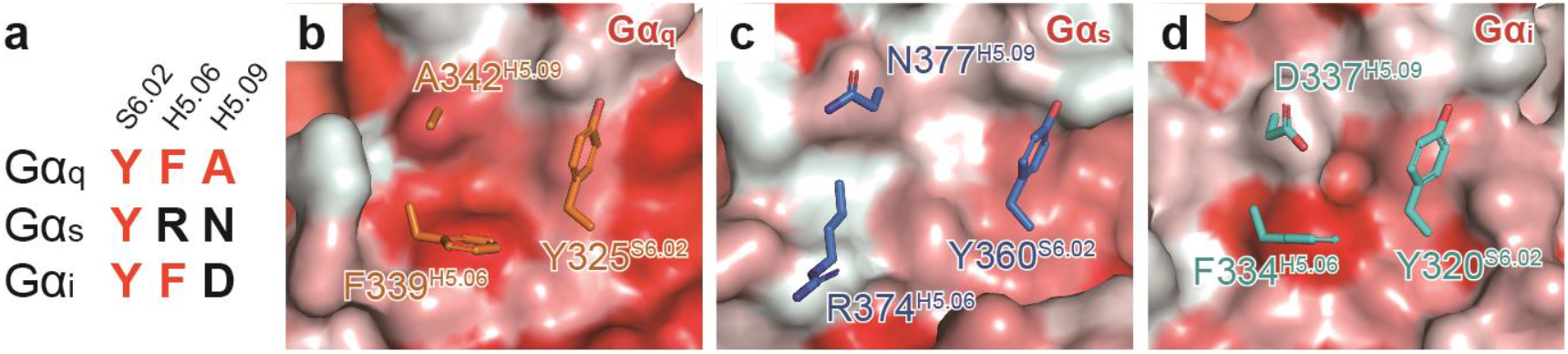
Comparison of the hydrophobic patch in Gα_q_ subunit to the corresponding sites in other G proteins. **a,** Sequence alignment of S6.02, H5.06, and H5.09 from Gα_q_, Gα_s_, and Gα_i_ subunits. Residues at positions S6.02, H5.06, and H5.09 comprise the hydrophobic patch to interact with CCK_A_R ICL3. **b-d**, Surface presentation of the patch by hydrophobicity. Side chains of residues at positions S6.02, H5.06, and H5.09 in Gα_q_ (**b**), Gα_s_ (**c**), and Gα_i_ (**d**) subunits are shown.

**Extended Data Table 1.**
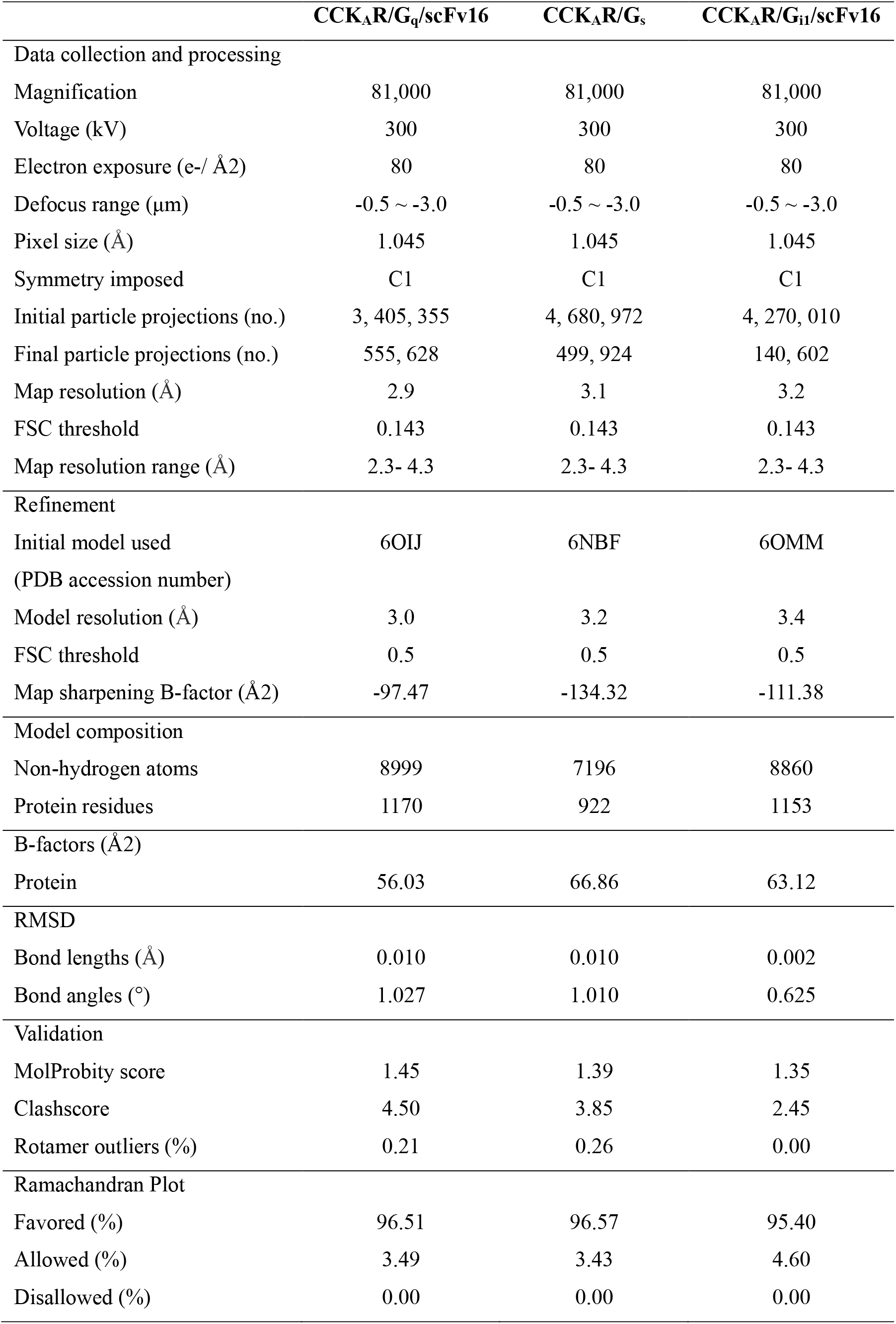
Cryo-EM data collection, refinement and validation statistics.

**Extended Data Table 2.** Effects of mutations in the ligand-binding pocket of CCK_A_R on CCK-8 binding affinities. Radiolabeled ligand ([^125I^]CCK-8) binding assay was performed to evaluate the ligand binding affinity of CCK_A_R mutants. Data represent mean pKi ± S.E.M. Experiments were performed in triplicate (n=3-4). \**P*<0.05 versus wild-type (WT). N.D., not determined. FACS analyses were performed to evaluate the surface expression of the CCK_A_R mutants.

| Mutant | pKi ± S.E.M. | Expression % |
| --- | --- | --- |
| WT | 8.58±0.12 | 100 |
| K105A | 7.78±0.22 | 77.86±6.24 |
| F107A | N.D. | 71.85±6.84 |
| T118A | 8.73±0.13 | 78.06±5.38 |
| M121A | 8.03±0.15 | 74.99±5.48 |
| V125A | 8.68±0.12 | 46.48±1.03 |
| Y176A | N.D. | 26.63±2.43 |
| F185A | 7.98±0.24 | 77.98±4.85 |
| M195A | 8.10±0.26 | 85.45±4.52 |
| C196A | N.D. | 3.24±0.06 |
| R197A | N.D. | 104.14±5.14 |
| H210A | 8.61±0.12 | 81.19±4.32 |
| I329A | N.D. | 74.22±7.37 |
| F330A | 8.67±0.09 | 27.31±2.74 |
| A332G | 8.43±0.12 | 32.50±4.21 |
| N333A | N.D. | 63.96±3.31 |
| R336A | N.D. | 82.96±5.35 |
| A343G | N.D. | 50.61±5.37 |
| E344A | N.D. | 88.00±13.77 |
| L347A | N.D. | 52.59±1.43 |
| S348A | N.D. | 98.35±8.18 |
| I352A | N.D. | 80.35±1.26 |
| Y360A | 8.00±0.09 | 82.85±6.85 |

**Extended Data Table 3.** Coupling activity of CCK_A_R with different G proteins. BRET assay was performed to evaluate the coupling activity of CCK_A_R with different G proteins. Data represent mean pEC_50_ ± S.E.M. Decreased fold of *E_max_* compared to G_q_ was calculated. Radiolabeled ligand binding assay was used to evaluate the allosteric effects of different G proteins on the binding affinity of CCK-8. The binding affinities are indicated as pKi ± S.E.M. All data were analyzed by two-tailed Student’s *t*-test. \**p*<0.05 versus receptor. Experiments were performed in triplicate (n=3).

| Group | pEC<sub>50</sub> ± S.E.M. | Decreased fold<br>of $E_{max}$ | pKi ± S.E.M. |
| --- | --- | --- | --- |
| Receptor | — | — | 7.92 ± 0.06 |
| Receptor + G<sub>q</sub> | 8.42 ± 0.08 | 1 | 8.28 ± 0.08* |
| Receptor + G<sub>i</sub> | 7.32 ± 0.22 | 6.60 | 7.87 ± 0.07 |
| Receptor + G<sub>s</sub> | 7.92 ± 0.65 | 20.33 | 8.02 ± 0.06 |

**Extended Data Table 4.** Effect of I296G mutation of CCK_A_R on G protein-coupling activity. BRET based NanoBiT G-protein recruitment and NanoBiT G-protein dissociation assays were performed to evaluate G_q_-, G_i_-, and G_s_-coupling activity, respectively. Data represent mean pEC_50_ ± S.E.M. \*\**p*<0.01 versus wild-type (WT). FACS analyses were performed to evaluate the surface expression of CCK_A_R mutant. Radiolabeled ligand binding assay was used to evaluate the effects of the mutation on the binding affinity of CCK-8. The binding affinities are indicated as pKi ± S.E.M. All data were analyzed by two-tailed Student’s *t*-test. Experiments were performed in triplicate (n=3).

| Mutant | pEC<sub>50</sub> ± S.E.M. |  |  | Expression % | pK<sub>i</sub> ± S.E.M. |
| --- | --- | --- | --- | --- | --- |
|  | G<sub>q</sub> | G<sub>i</sub> | G<sub>s</sub> |  |  |
| WT | 9.14 ± 0.04 | 6.81 ± 0.12 | 10.48 ± 0.10 | 100 | 8.58 ± 0.12 |
| I296G | 8.38 ± 0.09** | 6.66 ± 0.08 | 10.46 ± 0.20 | 99.52 ± 3.10 | 8.63 ± 0.13 |

